# Targeting Endothelial PERK Accelerates Lymphoid Regeneration by Enhancing DLL4-NOTCH3 Signaling at the Pre-B Niche

**DOI:** 10.1101/2025.10.15.682539

**Authors:** Bingqing Zou, Qiuyun Chen, Junjun Zheng, Yimin Ma, Jay Myers, Yinghui Shang, Mofei Huang, Paul Christensen, Stanley Adoro, Chih-hang Anthony Tang, Chih-Chi Andrew Hu, Sai Ravi Kiran Pingali, Wei Xin, Keith Syson Chan, Stephen Wong, Youli Zu, Hamed Jafar-Nejad, Lan Zhou

**Affiliations:** Department of Pathology and Genomic Medicine, Houston Methodist Research Institute, Houston, TX 77030, USA; Department of Pathology, Case Western Reserve University, Cleveland, OH 44106, USA; Center for Immunotherapy, Houston Methodist Neal Cancer Center, Houston Methodist Research Institute, Houston, TX 77030, USA; Department of Pediatrics, Case Western Reserve University, Cleveland, OH 44106, USA; Department of Hematology, Shanghai General Hospital, Shanghai Jiao Tong University School of Medicine, Shanghai 200080, China; Experimental Immunology Branch, National Cancer Institute, National Institutes of Health, Bethesda, MD 20892, USA; Houston Methodist Neal Cancer Center, Houston Methodist Research Institute Houston, TX 77030, USA; Department of Medicine, Houston Methodist Research Institute, Houston, TX 77030, USA; Department of Pathology, University of South Alabama, Mobile, AL 36688, USA; Department of Urology, Houston Methodist Research Institute, Houston, TX 77030, USA; Translational Biophotonics Laboratory, Department of Systems Medicine and Bioengineering, Houston Methodist Neal Cancer Center, Houston, TX 77030, USA; Department of Pathology and Laboratory Medicine and Microbiology and Immunology, Weill Medical College of Cornell University, New York City, NY 10021, USA; Department of Molecular & Human Genetics, Baylor College of Medicine, Houston, TX 77030, USA

## Abstract

Delayed immune recovery after hematopoietic stem cell (HSC) transplantation is associated with a poor clinical outcome, yet strategies to enhance lymphocyte regeneration are limited. We studied the role of unfolded protein response (ER stress) in hematopoietic regeneration within the bone marrow (BM) microenvironment. We revealed that PERK activation is a prominent feature of BM endothelium in leukemia patients and is a hallmark response in mouse BM following ionizing irradiation. Ablating endothelial *Perk* boosted Notch ligand DLL4 expression and promoted DLL4-dependent early HSC and B progenitor regeneration. Single-cell analysis shows that endothelial DLL4 activates NOTCH3 expressed by mesenchymal stroma cells, and that the PERK-DLL4 axis coordinates the regulation of lymphoid commitment and niche cytokine production. NOTCH3 is critical for the upregulation of IL7 following irradiation and for supporting the expansion of lymphoid progenitors in mesenchymal sphere cultures. These findings not only unveil a previously unrecognized ER stress-controlled vascular-stroma signaling mechanism in regenerative hematopoiesis but also highlight PERK blockade as a promising therapeutic strategy to improve immune recovery after myeloablative transplantation.

**Summary:** Zou et al unravel that the adaptive ER stress response in bone marrow blood vessels restricts the post-transplant regeneration of immune progenitor cells by attenuating the expression of Notch ligand DLL4. Targeting ER stress sensor PERK can accelerate immune recovery after transplantation by enhancing DLL4-NOTCH3 signaling and IL7 cytokine production.

## Introduction

Hematopoietic stem cells (HSCs) reside in a tightly controlled bone marrow (BM) niche that regulates and maintains hematopoietic homeostasis and regeneration (Morrison and Scadden 2014, Calvi and Link 2015, Pinho and Frenette 2019). Endothelial cells (ECs) and mesenchymal stroma cells (MSCs) (marked by LepR or Prx-1) provide critical pro-hematopoietic factors to support the HSC pool and IL7 for restricted progenitors including common lymphoid progenitors (CLPs) (Ding, Saunders et al. 2012, Ding and Morrison 2013, Greenbaum, Hsu et al. 2013, Cordeiro Gomes, Hara et al. 2016, Comazzetto, Murphy et al. 2019). Single cell and spatial transcriptomics analyses reveal that CXCL12-abundant reticular cells (CARs), which overlap with the Prx-1^+^ or LepR^+^ MSCs, show peri-vascular localization (Ding, Saunders et al. 2012, Cordeiro Gomes, Hara et al. 2016, Baccin, Al-Sabah et al. 2020). In addition, Notch ligands expressed by endothelium are known to function as an angiocrine factor regulating Notch-dependent HSC function (Butler, Nolan et al. 2010). Single cell transcriptome profiling revealed that Delta-like 4 (DLL4), a major Notch ligand expressed by BM ECs, displays dynamic regulation in response to 5-FU treatment (Tikhonova, Dolgalev et al. 2019), suggesting that regulating Notch signaling may be a key step in post-myeloablative hematopoietic regeneration.

A favorable outcome after HSC transplantation is dependent on timely HSC regeneration and differentiation into multiple blood cell lineages (Warr, Pietras et al. 2011). In hematopoietic regeneration, the unfolded protein response (UPR), which is an adaptive response to endoplasmic reticulum (ER) stress, is critical for meeting the increased demand for protein synthesis required for rapid cellular proliferation. ER stress response also helps adaptation to conditions such as increased levels of reactive oxygen species (ROS) generated by ionizing radiation and chemotherapy (van Galen, Kreso et al. 2014). ROS may cause further damage to the macromolecules leading to protein misfolding and unfolding (Mikkelsen and Wardman 2003, Malhotra and Kaufman 2007, Ding, Yang et al. 2012). As part of UPR, activated PKR-Like ER Kinase (PERK; encoded by *Eif2ak3*) phosphorylates the alpha subunit of eukaryotic initiation factor-2 (eIF-2α), which in turn inhibits general translation initiation to restore ER homeostasis. Our recent work revealed that endothelial PERK plays an active role in the regulation of Notch ligand expression and leukemia-induced vascular niche remodeling (Liu, Chen et al. 2022).

However, how ER stress response in the BM stroma adapts to myeloablation and impacts hematopoietic stem cell and progenitor (HSPC) recovery remains unclear. To address the role of ER stress response in HSPC regeneration, we performed BM transplantation, imaging, and MSC functional analysis. We revealed that perturbation of endothelial *Perk* boosted DLL4 expression and promoted DLL4-dependent HSPC regeneration. Importantly, EC-expressed DLL4 and stroma-expressed NOTCH3 co-regulate lymphoid priming of regenerative HSC. Furthermore, we used single-cell analysis to define the cellular and molecular networks required for enhanced lymphoid recovery by targeting endothelial PERK-DLL4 axis, highlighting a previously unrecognized vascular-stroma signaling mechanism in regenerative hematopoiesis.

## Results

### Ablating endothelial PERK or blocking PERK activation promotes early HSC and lymphoid regeneration

We investigated whether PERK is activated in the BM microenvironment during transplantation. Forty-eight hours after lethal irradiation, the level of phosphor-PERK (p-PERK) was markedly increased in the BM ECs but not in MSCs (Fig 1A). This selective activation prompted us to focus on endothelial PERK under irradiation-induced stress hematopoiesis. We generated mice that have PERK deleted in ECs by tamoxifen (*VE-cadherin^ERT2-Cre^/Perk^F/F^*or *Perk*^iΔEC^ mice). Analysis of recovering BM CD31^+^ ECs of *Perk*^iΔEC^ mice revealed successful ablation of PERK activation (Fig 1B). Using this mouse model, we investigated the impact of PERK ablation on hematopoietic recovery following transplantation and chemotherapy. We infused wild type (WT) BM cells (Ly5.1) into irradiated *Perk*^iΔEC^ mice (Ly5.2) or littermate controls (*VE-cadherin^ERT2-Cre^/PERK^F/+^* or *PERK^F/F^*). White blood cell (WBC) counts consistently elevated by 17∼42% over 1∼4 months post-transplantation in *Perk*^iΔEC^ mice. The most pronounced effect was increase in donor-derived B lymphocytes in *Perk*^iΔEC^ mice compared to controls (Fig 1C). The absolute numbers of HSC, CLP, and B lineage progenitors (pre-proB, pro-B, and pre-B cells) all increased at 1-month in transplanted *Perk*^iΔEC^ mice (Fig 1D; supplementary Fig 1A). By 4 months, there were no significant differences in the numbers of HSC/CLP/B progenitors in *Perk*^iΔEC^ mice compared to controls (Fig 1E). We then assessed hematopoietic recovery after 5-FU treatment. *Perk*^iΔEC^ mice initially displayed mitigation of WBC decline. WBC continued to recover faster on day ten and day twenty, with elevated B cell numbers and decreased myeloid output (Fig 1F). To explore the regenerative potential of targeting PERK, we treated mice transplanted with WT BM cells with a PERK inhibitor (PERKi) and performed hematopoietic analysis on day 21, a clinically relevant time point for assessing early lymphocyte recovery (Kim, Kim et al. 2004). PERKi treatment increased circulating B cells (Fig 1H) and HSC/lymphoid progenitor regeneration (Fig 1I-J). Blocking IRE1α/XBP1, the other major arm of ER stress activation (Ranatunga, Tang et al. 2014, Tang, Ranatunga et al. 2014) did not affect lymphoid recovery or progenitor regeneration (supplementary Fig 1B-D), indicating this is a PERK-dependent effect. Consistently, we observed mild increase in the expression of the spliced form of XBP1 after lethal irradiation (data not shown). Therefore, PERK blockade or genetically ablating endothelial PERK mitigated myelosuppression but enhanced lymphoid recovery in transplantation without impacting long term hematopoiesis.

**Figure 1.**
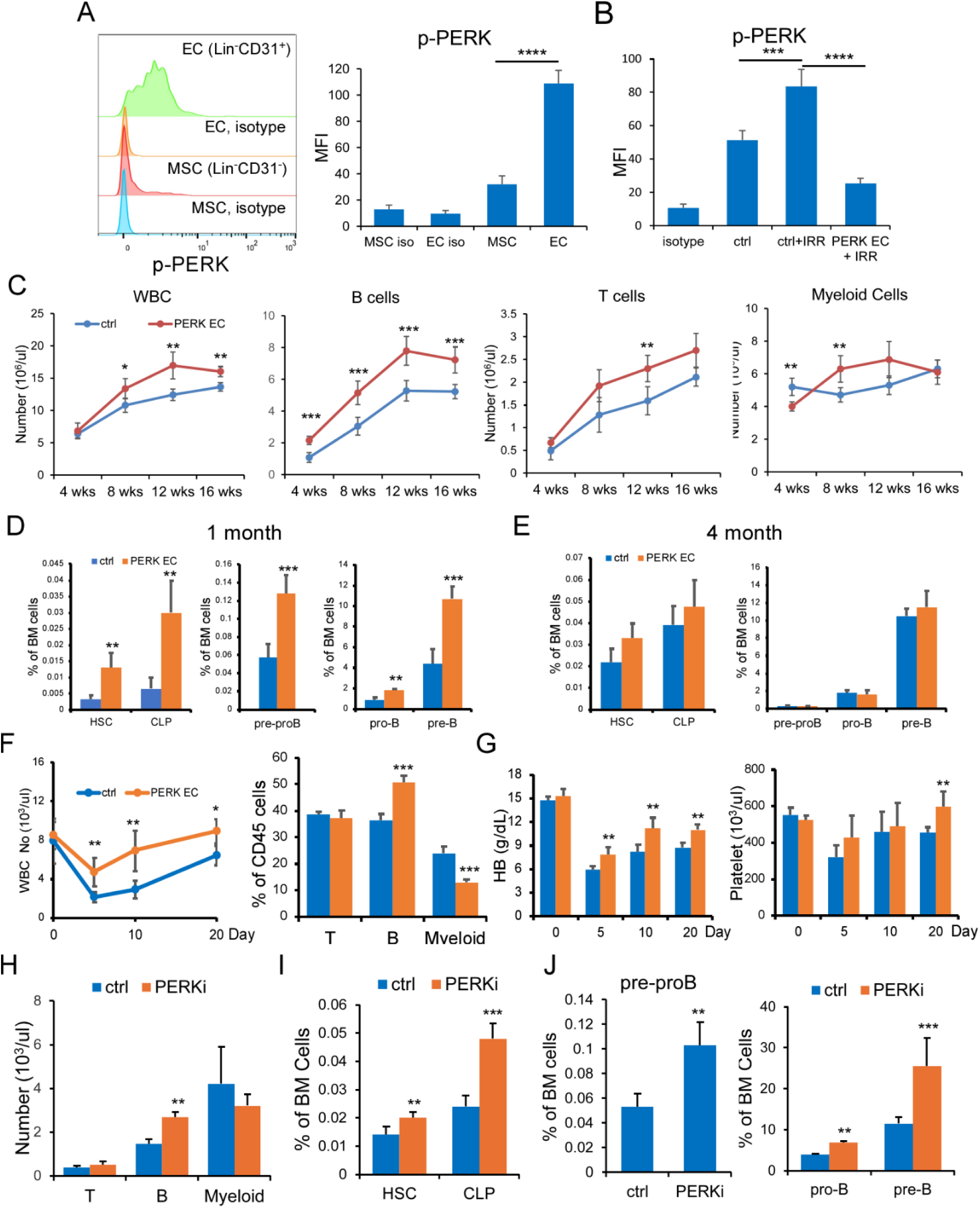
Ablating endothelial PERK promotes early HSC and lymphoid progenitor regeneration. (A) FACS analysis was caried out on MACSQuant Analyzer 16. Representative FACS profile and mean fluorescent intensity (MFI) of activated PERK (p-PERK) from 3 similar experiments in bone marrow endothelial cells (ECs) and mesenchymal stroma cells (MSCs). MSC is defined by Lin (CD11b, Gr1, B220, CD4, CD8, Ter119, NK1.1)^-^ CD31^-^ while EC is defined by Lin^-^CD31^+^. (B) MFI of p-PERK in non-irradiated wild type (WT) (ctrl), irradiated WT (ctrl + IRR) and irradiated *Perk*^iΔEC^ mice (PERK EC + IRR) at 48 hours following 850 cGy (Cesium-137) irradiation. MFI of p-PERK in A-B is presented as averages ± SEM (n=4-5/each condition). (C) Peripheral blood was analyzed for WBC counts, B cells, T cells, and granulocytes at 4, 8, 12, and 16 weeks after transplantation (n=8-9/group; data pooled from 3 experiments). (D) The frequencies of HSC (Lin^−^Sca-1^+^c-Kit^+^CD150^+^CD48^−^CD34^-^Flk2^−^), CLP (Lin^−^IL7R^+^Flt3^+^Sca-1^low^c-Kit^+^low) and B progenitors (pre-proB: Lin [CD11b, Gr1, CD3, NK1.1, Ter119)^-^B220^+^CD43^+^Flt3^+^CD19^-^; pro-B: Lin^-^B220^+^CD43^+^Flt3^-^CD19^+^; pre-B: Lin^-^B220^+^CD43^-^) were determined in control and *Perk*^iΔEC^ mice BM at 1 month after transplantation (n=6-7/group; data pooled from 2 experiments). (E) The frequencies of HSC, CLP, and B progenitors were determined in control or *Perk*^iΔEC^ mice at 4 months after transplantation (n=8-9/group; data pooled from 3 experiments). (F) WBC counts were determined on days 5, 10, and 20 after single dose 5-FU treatment. Frequencies and absolute numbers (not shown) of B cells, T cells, and granulocytes were determined on day 20. Absolute numbers showed similar changes. (G) Hemoglobin levels and platelet counts were determined on day 20. Results (F-G) are presented as averages ± SEM (n=5-6/group). (H) WT mice transplanted with WT BM cells received PERK inhibitor (PERKi; GSK2656157) (50 mg/kg) or control treatment twice daily for 2 weeks starting on day 3 after transplantation. Peripheral blood B, T and granulocytes were enumerated on day 21. (I-J) BM HSC, CLP (I) and B progenitors (J) were also determined. Results were pooled from 3 experiments and presented as averages ± SEM (n=9-12/ group). \**P*< 0.05, \*\**P*<0.01, *** *P*<0.001.

### DLL4 regulated by PERK is indispensable for post-irradiation HSC and lymphoid progenitor recovery

DLL4 is the major Notch ligand expressed by BM ECs and is downregulated after chemotherapy (Tikhonova, Dolgalev et al. 2019), but the mechanism of this downregulation was unknown. We found that DLL4 level largely unchanged in ECs after irradiation (Fig 2A). Intriguingly, its expression increased by 60% in PERK-deleted ECs (Fig 2A-B). Blocking PERK suppresses activation of eIF-2α (p-EIF2α) and upregulates DLL4 protein expression in irradiated BM ECs *in vitro* (supplementary Fig 2A), unveiling a link between PERK and DLL4 regulation. To assess if DLL4 and PERK expression show similar changes in response to myeloablation in human disease, we evaluated BM specimens from acute myeloid leukemia (AML) patients who underwent myeloablative chemotherapy. We observed disorganized vessels and decreased cytoplasmic/membrane staining of DLL4 in AML BM when compared to tissues from individuals without hematologic malignancies. In contrast, expression of p-PERK enhanced in AML BM vessels (Fig 2C-D). To distinguish if enhanced HSC and lymphoid recovery in *Perk*^iΔEC^ mice is regulated by DLL4, we generated endothelial *Dll4* and *Perk* double knockout (DKO) mice (*VE-cadherin^ERT2-Cre^/Perk^F/F^*/*Dll4^F/F^*) and confirmed the complete extinction of DLL4 in DKO mice ECs (Fig 2A-B). We performed BM transplantation in DKO mice. B cells decreased while granulocytes increased in DKO recipients 1 month after receiving WT cells (Fig 2E). Unlike *Perk*^iΔEC^ recipients, which had increased HSCs and CLPs, DKO mice displayed decreased HSC and CLP recovery (Fig 2F). B progenitors reduced markedly (Fig 2G) while myeloid progenitors showed no significant changes (supplementary Fig 2B). Therefore, ablating *Dll4* completely abrogated the early increase of HSC and B progenitors mediated by *Perk* deletion.

**Figure 2.**
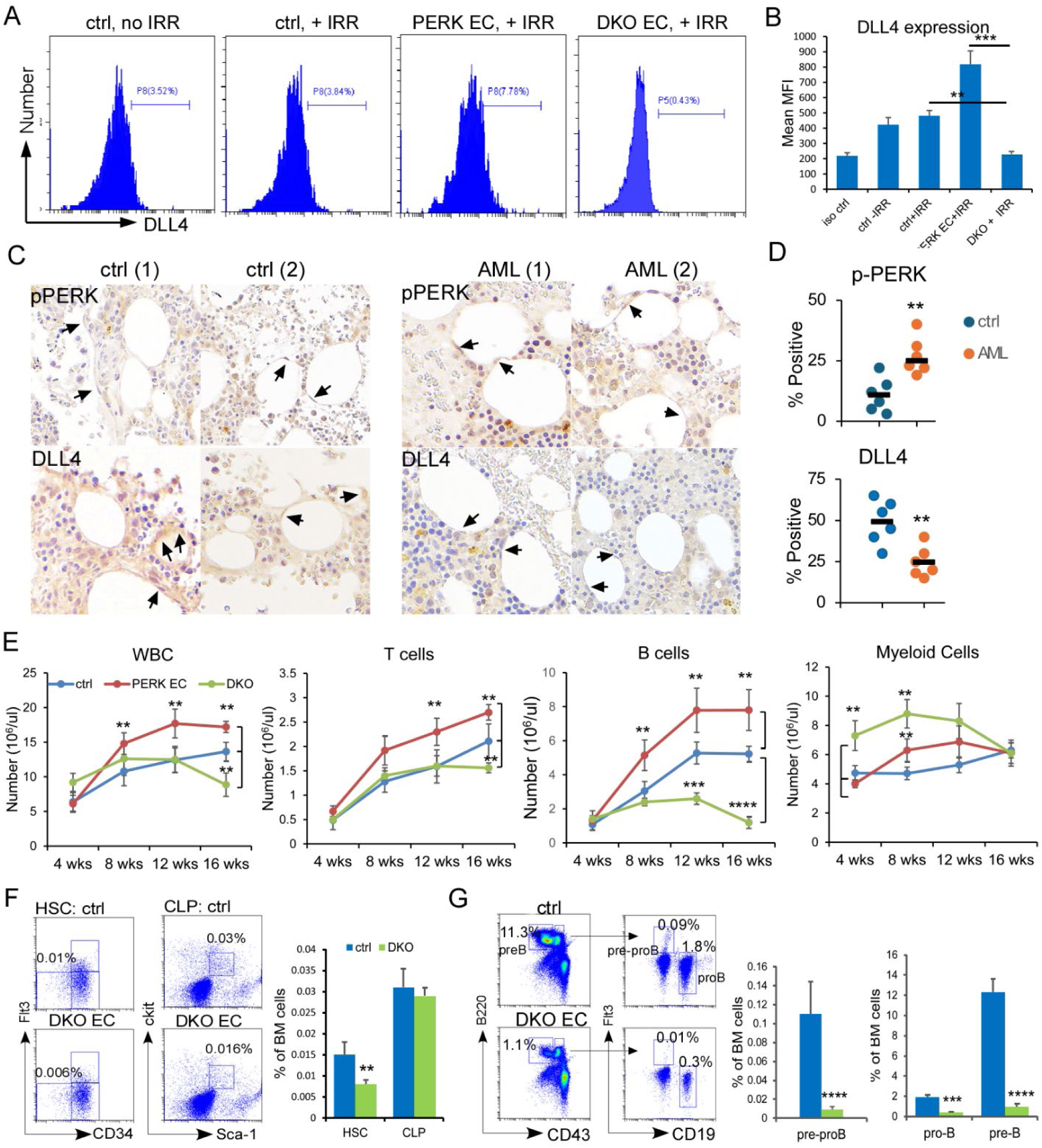
Endothelial DLL4 is regulated by PERK and is indispensable for post-irradiation HSC and B lymphoid progenitor recovery. (A) FACS analysis was carried out on CytoFLEX. Representative FACS profile from 3 similar experiments of BM endothelial DLL4 (Lin^-^ CD31/CD144^+^) expression at 72 hours in non-irradiated WT mice (ctrl, - IRR), WT irradiated mice (ctrl, + IRR), *Perk*^iΔEC^ irradiated mice (PERK EC, +IRR), and DKO irradiated mice (DKO + IRR) following 850 cGy irradiation. (B) MFI of DLL4 expression is presented as averages ± SEM (n=4-6/group). (C-D) Representative IHC staining images of endothelial DLL4 and p-PERK (arrows) (C) and percentage of positive cells (count ≥ 20 cells each case) (D) in human BM tissues from 6 non-neoplastic (ctrl) and 6 AML cases. Images were scanned and analyzed on MoticEasyScan Pro 6 (Motic) under 40 x magnification. (E) Enumeration of WBC and FACS analysis of B cells, T cells, and granulocytes at 4, 8, 12, and 16 weeks after transplantation (n=9-10/group; pooled from 3 experiments). (F-G) Representative FACS profile and frequencies of HSC/CLP (F) and B progenitors (G) in control and DKO mice at 1 month after transplantation. Results (E-G) are presented as averages ± SEM (n= 9-10/group). \*\**P*<0.01, *** *P*<0.001, **** *P*<0.0001

We then found that ablating EC *Dll4* alone (*VE-cadherin^ERT2-Cre^/DLL4^F/F^*, or *Dll4*^iΔEC^) is sufficient to impair HSPC regeneration. Like DKO mice, *Dll4*^iΔEC^ mice transplanted with WT BM displayed increased granulocytes but decreased B cells, HSCs, and CLPs (Supplementary Fig 2C-D). B progenitors were variably depleted in *Dll4*^iΔEC^ mice while GMPs expanded (supplementary Fig 2E-F). Notably, suppression of HSCs and B progenitors persisted 4 months after transplantation while GMPs normalized in *Dll4*^iΔEC^ mice by this age (supplemental Fig 2G-I). Considered together, these findings suggest that endothelial DLL4 in post-irradiation BM is required for HSPC and B lymphoid regeneration. The data suggests that in this context *Dll4* is epistatic to *Perk*, and its upregulation mediates the enhanced regeneration of HSCs and B progenitors observed upon *Perk* ablation.

### PERK-DLL4 regulates regenerative vascular homeostasis and integrity

DLL4 has an important role in angiogenesis (Benedito, Roca et al. 2009). We investigated the impact of endothelial PERK-DLL4 axis on post-irradiation vascular regeneration. To mitigate the interference from transplanted BM cells, we performed non-lethal irradiation (550 cGy) to induce vascular damage and regeneration. It has been shown that the collapsed vascular network rejuvenates by 2 weeks after whole body irradiation (Termini, Pang et al. 2021). Total marrow ECs decreased in post-irradiated *Dll4*^iΔEC^ mice but the EC number was not significantly reduced in irradiated *Perk*^iΔEC^ mice (Fig 3A-B). However, confocal analysis showed expansion of the vascular volume and increased Sca-1^+^ arterioles in the BM of *Dll4*^iΔEC^ mice (Fig 3C). This discrepancy is likely caused by decreased EC survival as we observed that more CD31^+^ cells of *Dll4*^iΔEC^ mice were labeled by DAPI than control ECs (not shown). In addition, the BM vessel bed in *Dll4*^iΔEC^ mice displayed a distorted pattern characterized by reduced vessel lengths and disrupted structures, suggesting that DLL4 may restrict post-irradiation disorganized angiogenesis.

**Figure 3.**
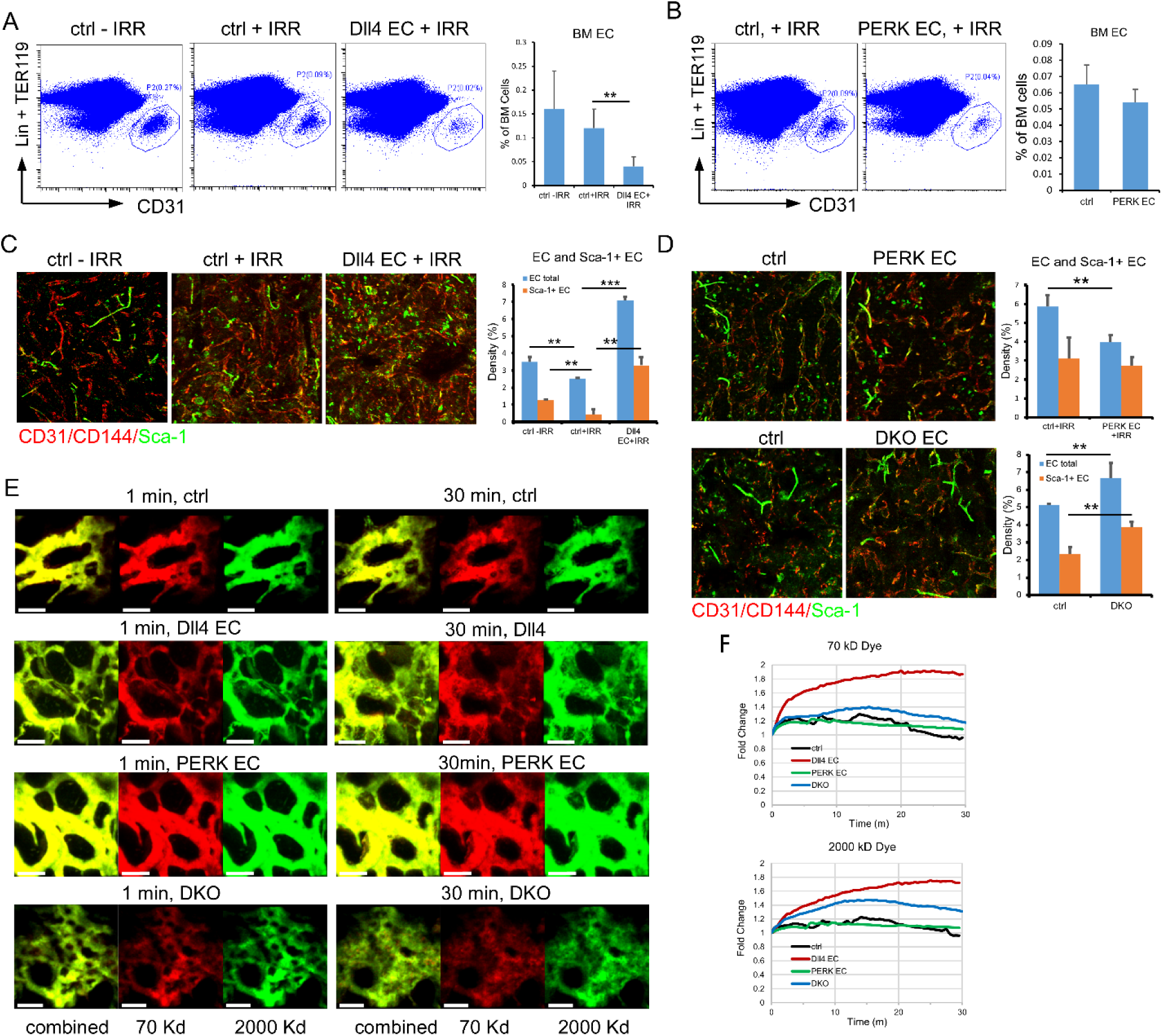
PERK-DLL4 regulates regenerative vascular homeostasis and integrity. (A-B) FACS analysis was carried out on CytoFLEX. Representative FACS profile of BM CD31^+^ ECs (Lin^-^CD31^+^) and frequencies of ECs in non-irradiated WT mice (ctrl, - IRR), in 550 cGy irradiated WT mice (ctrl, + IRR), 550 cGy irradiated *Dll4*^iΔEC^ mice (DLL4 EC, +IRR) (A), and 550 cGy irradiated *Perk*^iΔEC^ mice (PERK EC, +IRR) (B). (C-D) Whole-mount immunostaining of BM vascular bed with Alexa Fluor 647-anti-CD31, Alexa Fluor 647-anti-CD144 and FITC-anti-Sca-1 in WT and *Dll4*^iΔEC^ mice (C), WT, *Perk*^iΔEC^ and DKO mice (D). Representative images were generated in Imaris based on the corresponding fluorescent signals (left). The percentage of total CD31^+^/CD144^+^ volume (EC total) and CD31^+^Sca-1^+^ arteriole volume (Sca-1^+^ EC) were determined (right). (E) Intravital 2-photon imaging of calvarium BM vessels was performed in 3 independent experiments to determine vessel leakiness after retro-orbital injection of 2000 kD FITC-Dextran and 70 kD TRITC-Dextran immediately before imaging. Three dimensional images consisting of XY: 775 mm x 775 mm and Z: 140 mm (5 mm/step), were obtained every 30 seconds for 30 minutes. Images were drift-corrected, smoothed, and surfaces were generated on both FITC and TRITC signals. Representative images taken at 1min and 30 min were shown. (F) The corresponding total volumes for each signal were calculated at each timepoint and normalized to the first timepoint. Results in bar graphs (A-D) are presented as mean ± S.D (n=5-6/group). \*\**P*<0.01, \*\*\**P*<0.001.

In contrast, irradiated *Perk*^iΔEC^ mice, which expressed increased DLL4, showed the opposite phenotype, with a more organized BM vasculature (Fig 3B, upper panel). While visual inspection of DKO marrow vasculature might exhibit less disruption compared to the *Dll4*^iΔEC^ mice (Fig 3D, lower left panels), EC density quantification demonstrated an increase in DKO BM compared to control BM (Fig 3D, lower right panel), suggesting again that *Dll4* functions downstream of *Perk* in this context. To investigate if the PERK-DLL4 axis regulates vascular integrity, we monitored BM vessel leakage at 2 weeks after 550 cGy irradiation by intravital microscopy using both low- (LMW; 70 KD) and high molecular weight Dextran (HMW; 2000 KD). DLL4-deficient vessels were markedly more permeable to both LMW and HMW Dextran tracers (Fig 3E-F), indicating compromised barrier function. In comparison, *Perk* ablation alone had no significant effect on vascular permeability while DKO mice showed increased leakiness compared to *Perk*^iΔEC^ mice (Fig 3E-F). These results establish DLL4 as a critical determinant of vascular integrity following irradiation. Further, elevated DLL4 expression upon PERK ablation restricts abnormal angiogenesis and stabilizes the BM vasculature, thereby mitigating irradiation-induced vascular niche disruption.

### PERK and DLL4 differentially regulate pathways supporting B progenitor and HSC regeneration

To elucidate how endothelial PERK and DLL4 differentially regulate HSC and B progenitor regeneration, we performed scRNA-seq of regenerating BM cells on day eleven after sublethal irradiation. Under this condition, we were able to study regenerative BM without interference from infused donor cells. We identified fourteen cell populations based on canonical markers (Fig 4A; supplementary Fig 3A). Consistent with the transplant model, *Dll4*^iΔEC^ mice displayed an expansion of myeloid progenitors (GMPs) and a reduction of B progenitors (preB/B), while DKO mice exhibited a profound loss of preB/B cells. In contrast, *Perk*^iΔEC^ mice showed a decrease in GMPs accompanied by an increase in B progenitors (Fig 4B). Pathway analysis in B progenitors revealed that defense response and humoral immune response were upregulated upon *Perk* extinction but downregulated with *Dll4* ablation (Fig 4C). This was supported by the upregulation of genes implicated in lymphoid development and function in B progenitors of *Perk*^iΔEC^ mice (Fig 4D). In line with increased B progenitor production, *Il7* and *Il7r* expression increased in *Perk*^iΔEC^ stroma cells and HSPC cells, respectively, but decreased in *Dll4*^iΔEC^ and DKO mice (Fig 4E).

**Figure 4.**
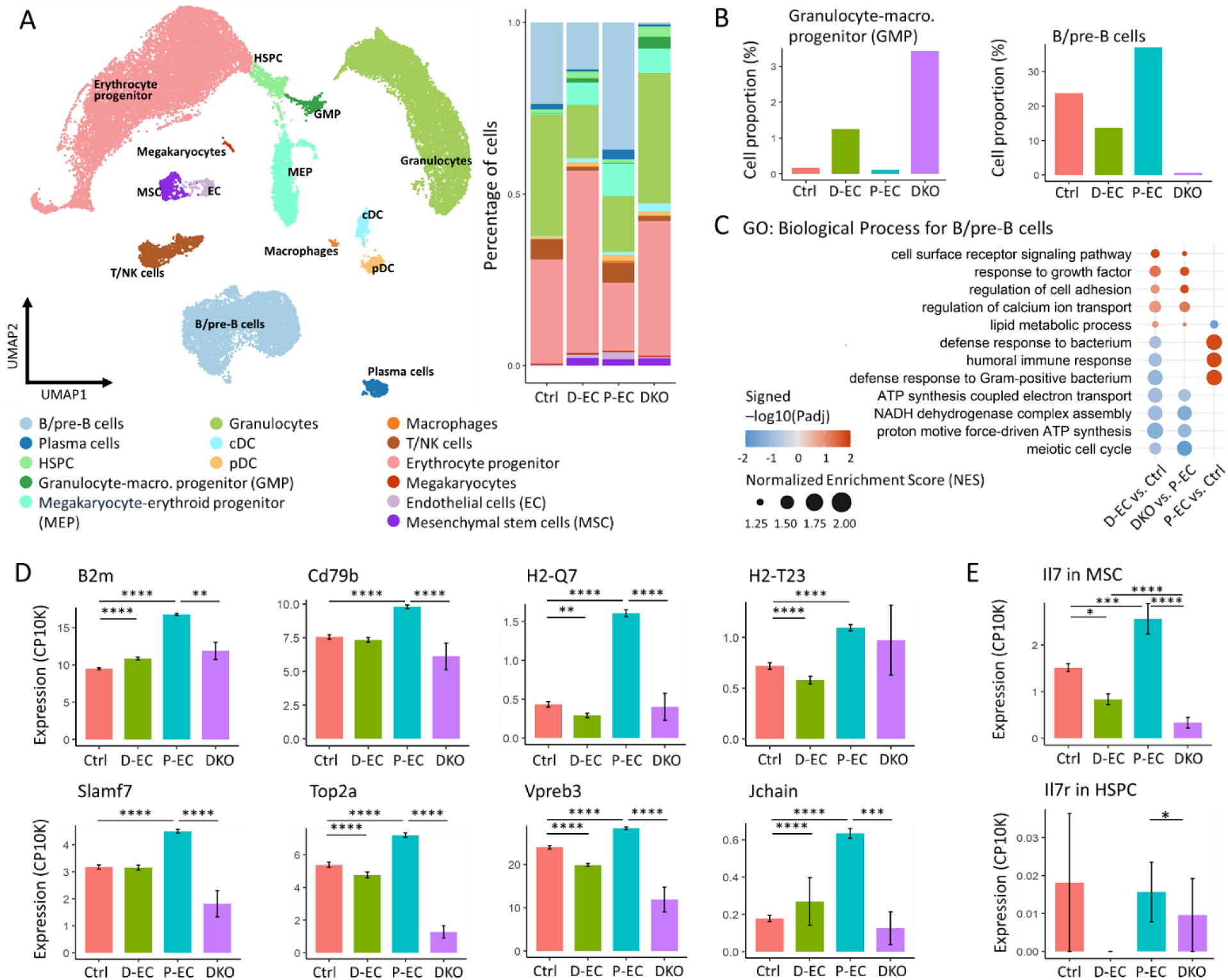
ScRNA-seq reveals BM cellular and molecular network supporting B progenitor development. (A) UMAP visualization and proportion of identified cell clusters from scRNA-seq analysis in 4 groups of mice: control (ctrl) (n=4), *Dll4*^iΔEC^ (D-EC) (n=4), *Perk*^iΔEC^ (P-EC) (n=4) and DKO (n=3). (B) Proportions of GMP and B/pre-B cells in four groups of mice. (C) Integrated dot plots for GSEA results in Gene Ontology-Biological Processes (GO-BP) pathway, illustrating significant changes identified when comparing *Dll4*^iΔEC^ mice vs control (D-EC vs Ctrl), *Perk*^iΔEC^ mice vs control (P-EC vs Ctrl), and DKO vs *Perk*^iΔEC^ (DKO vs PP-EC) for B/pre-B cells. Dot sizes demonstrated the normalized enrichment scores (NES) and color codes visualized significance where red indicates upregulated pathways and blue indicates down-regulated pathways. (D) Differentially expressed genes for B/pre-B cells. Statistical significances were first determined by Kruskal–Wallis test across four conditions, followed by post-hoc pairwise tests using MAST algorithm. (E) *Il7* gene expression in MSC cells and *Il7r* gene expression in HSPC cells. Statistical significances were first determined by Kruskal–Wallis test across four conditions (adjusted P<0.05), followed by post-hoc pairwise tests using MAST algorithm. * *P* < 0.05, ** *P* < 0.01, *** *P* < 0.001, **** *P* <0.0001.

Similarly, in HSPCs, immune response and defensive response were upregulated by *Perk* ablation and downregulated with *Dll4* ablation (Fig 5A). Analysis of the top differentially regulated genes in emerging HSPCs (Fig 5B) showed upregulation of the B-cell specific activating transcription factor (TF) *Pax5* and HSC marker genes (*Ly6a, Adgrl4*) (Spangrude, Heimfeld et al. 1988, Solaimani Kartalaei, Yamada-Inagawa et al. 2015) in *Perk*^iΔEC^ mice, but downregulation of these genes by *Dll4* ablation (Fig 5C). Genes regulating HSC quiescence (*Cd53* and *Emb*) (Silberstein, Goncalves et al. 2016, Greenberg, Paracatu et al. 2023) and self-renewal or regeneration (*Mecom and Stat3*)(Sato, Kamio et al. 2020, Voit, Tao et al. 2023, Patel, Zhou et al. 2024) were also upregulated in *Perk*^iΔEC^ mice but downregulated or unchanged in *Dll4*^iΔEC^ and DKO mice. The downregulation of genes supporting HSC quiescence in *Dll4*^iΔEC^ mice prompted us to assess the role of *Dll4* in HSC regeneration. We transplanted WT BM cells (Ly5.1) into lethally irradiated *Dll4*^iΔEC^ and control mice (Ly5.2) to reconstitute HSC pools (Fig 5D). Cell cycle analysis 1 month after transplantation showed that donor-derived HSCs in *Dll4*^iΔEC^ mice were cycling faster than those in control mice leading to a reduction of quiescent HSCs in the G_0_ phase (Fig 5E). We isolated BM cells from the primary WT or *Dll4*^iΔEC^ recipients 4 months after transplant, normalized for HSC frequency with competitor BM cells (Ly5.2), and performed competitive secondary and tertiary transplantations (Fig 5D). The engraftment of donors pre-conditioned in *Dll4*^iΔEC^ mice significantly reduced in both secondary and tertiary *Dll4*^iΔEC^ recipients when compared to cells pre-conditioned in WT mice and engrafted in WT recipients (Fig 5F). Consistently, fewer HSPCs isolated from secondary competitive transplant *Dll4*^iΔEC^ recipient mice were in G_0_ phase (Fig 5G). Collectively, these findings demonstrate that endothelial DLL4 is not only essential for HSC and B progenitor regeneration but also plays a key role in HSC quiescence maintenance which is essential for long-term hematopoietic regeneration.

**Figure 5.**
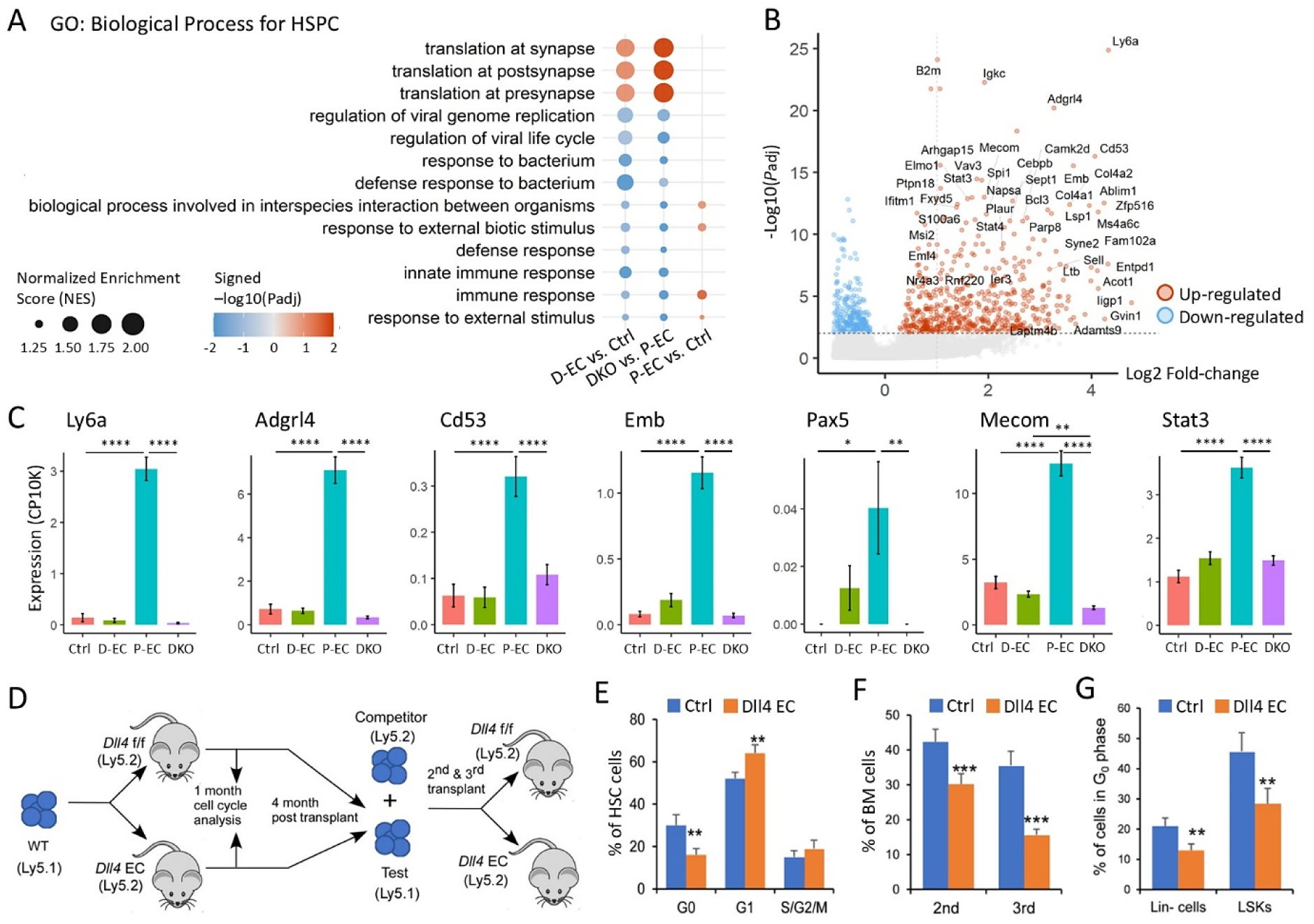
PERK and DLL4 differentially regulate molecular pathways supporting HSC regeneration. (A) Integrated dot plots for GSEA results in Gene Ontology-Biological Processes (GO-BP) pathway, illustrating significant changes identified when comparing *Dll4*^iΔEC^ mice vs control (D-EC vs Ctrl), *Perk*^iΔEC^ mice vs control (P-EC vs Ctrl), and DKO vs *Perk*^iΔEC^ (DKO vs PP-EC) for HSPC cells. Dot sizes demonstrated the NES and color codes visualized significance where red indicates upregulated pathways and blue indicates down-regulated pathways. (B) Volcano plot shows DEGs of *Perk*^iΔEC^ (P-EC) vs. CTRL derived from MAST algorithm. (C) Differentially expressed genes for HSPC cells. Statistical significances were first determined by Kruskal–Wallis test across four conditions, followed by post-hoc pairwise tests using MAST algorithm. * *P* < 0.05, ** *P* < 0.01, *** *P* < 0.001, **** *P* <0.0001. (D) Scheme of primary BM transplantation and competitive 2^nd^ and 3^rd^ transplantation. (E) Marrow cells were stained with lineage antibodies (CD4, CD8, B220, Gr-1, CD11b, TER119, and NK1.1), c-kit, Sca-1, CD34, Flt3, CD150, pyronin Y (RNA dye), and Hoechst 33342 (DNA dye). The relative proportion of cells in G_0_, G_1_, and S/G_2_/M phase of the cell cycle was analyzed on gated HSCs (Ly5.1) (n=4-6/group). (F) Proportion of control and *Dll4*^iΔEC^ primed donor-derived BM cells (Ly5.1) in 2^nd^ or 3^rd^ transplant recipients. (G) Proportion of Lin- and LSK cells (Ly5.1) in G_0_ phase in 2^nd^ transplant recipients. Results in bar graphs (E-G) are presented as mean ± SEM. \*\**P*<0.01, \*\*\**P*<0.001.

### PERK-DLL4 axis orchestrates the stroma network for HSC lymphoid priming

We further examined signaling pathways and HSC-promoting cytokines in the regenerative stroma. We distinguished four major cell clusters, including CARs, osteoblasts, sinusoidal ECs and arterioles (Fig 6A). CARs displayed the highest expression levels of CXCL12 and SCF (KITL) and were separated into two subsets: the adipogenic lineage precursors (MALP) (Zhong, Yao et al. 2020) or adipo-CAR cells and the osteogenic CAR or osteo-CAR (supplemental Fig 3B) (Baccin, Al-Sabah et al. 2020). Pathway analysis revealed an enhanced response to IFNβ and pyroptosis in *Perk*^iΔEC^ ECs (Fig 6B). We then examined HSC cytokine gene expression. *Il7* expression increased 2-fold in adipo-CARs of *Perk*^iΔEC^ mice, its expression markedly decreased in *Dll4*^iΔEC^ and DKO mice (Fig 6C). *Cxcl12 and Kitl* displayed a milder but significant increase and decrease in *Perk*^iΔEC^ and DKO mice, respectively (supplemental Fig 4A). We queried the influences of these HSC supporting cytokines on various cellular compartments by CellChat analysis. *Il7* from adipo- and osteo-CARs displayed strong communication with *Il7r* present on preB/B, T/NK, and MEP cells in *Perk*^iΔEC^ mice, but these interactions were much weaker in *Dll4*^iΔEC^ or DKO mice (supplementary Fig 4B). CXCL12 from CARs and arterioles strongly interacted with CXCR4 in preB/B cells in *Perk*^iΔEC^ mice while it showed the strongest communication with MEPs in DLL4-deficient mice. There were substantial interactions between SCF (*Kitl*) in arteriolar ECs and CARs with HSPC and GMPs in control BM. These interactions were enhanced in *Perk*^iΔEC^ mice but repressed in *Dll4*^iΔEC^ and DKO mice (supplementary Fig 4C-D).

**Figure 6.**
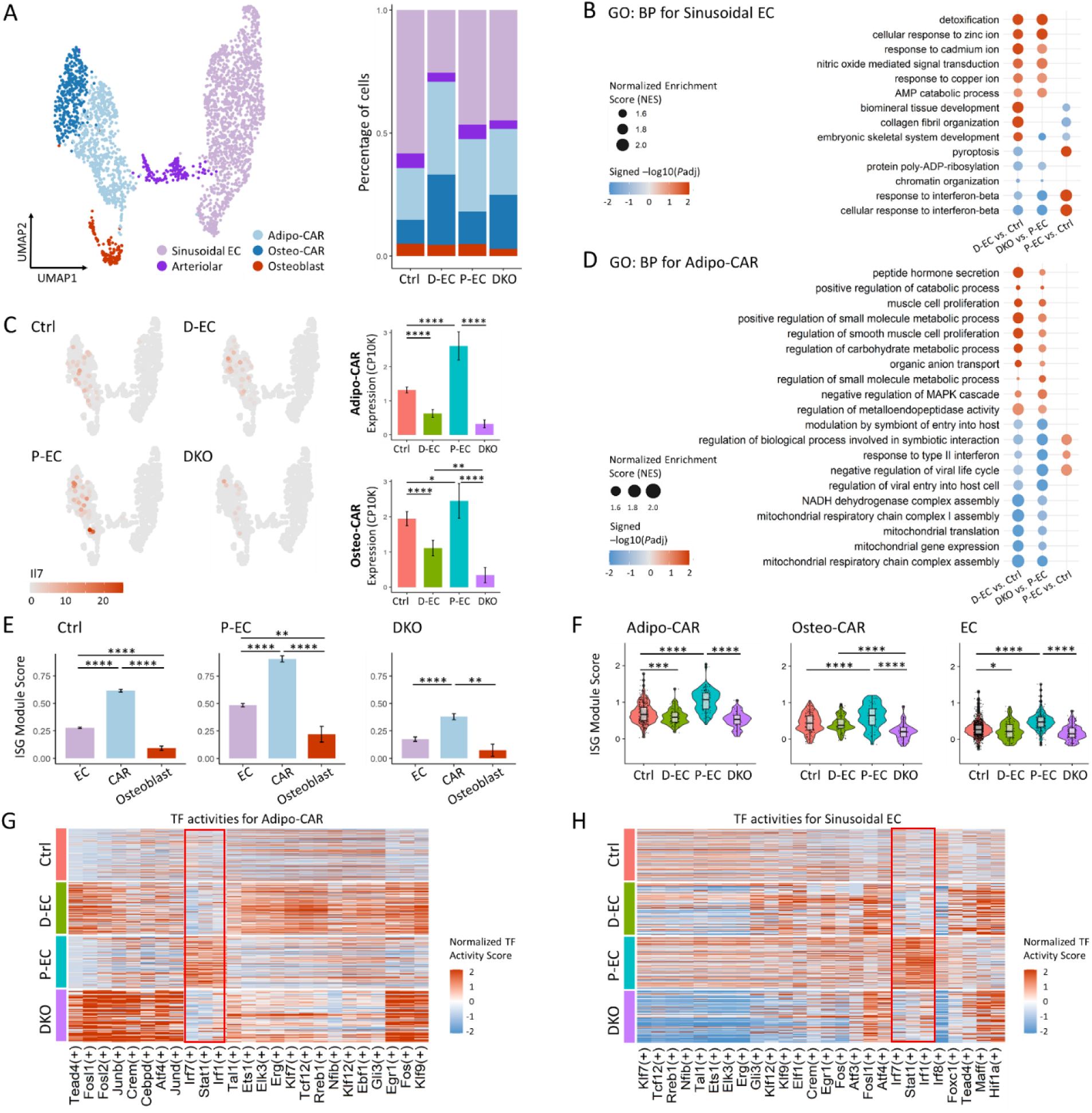
Signaling pathways and HSPC-promoting cytokines in the regenerative stroma network regulated by PERK-DLL4 axis. (A) UMAP visualization and proportion of five stromal cell subsets from scRNA-seq analysis. (B) Integrated dot plots for significant GO-BP pathway, illustrating significant changes identified when comparing *Dll4*^iΔEC^ mice vs control (D-EC vs Ctrl), *Perk*^iΔEC^ mice vs control (P-EC vs Ctrl), and DKO vs *Perk*^iΔEC^ (DKO vs PP-EC) for sinusoidal ECs. Dot sizes demonstrated the NES and color codes visualized significance where red indicates upregulated pathways and blue indicates down-regulated pathways. (C) UMAP and bar plots of *Il7* expression across four groups of mice. Statistical significances were first determined by Kruskal–Wallis test across four conditions, followed by post-hoc pairwise tests using MAST algorithm. * *P* < 0.05, ** *P* < 0.01, *** *P* < 0.001, **** *P* <0.0001. (D) Integrated dot plots for significant GO-BP pathway, illustrating significant changes identified when comparing *Dll4*^iΔEC^ mice vs control (D-EC vs Ctrl), *Perk*^iΔEC^ mice vs control (P-EC vs Ctrl), and DKO vs *Perk*^iΔEC^ (DKO vs PP-EC) for adipo-CAR cells. Dot sizes demonstrate NES and color codes visualized significance where red indicates upregulated and blue down-regulated pathways. (E) Interferon-Stimulated Gene (ISG) score defined by BP pathway (*Ifi35, Ifi44, Ifit1, Ifi73, Il7, Irf7, Oas3, Bst2, Cxcl10, Cxcl12, Ddx60, Hsd17b1, and Stat1*) in stroma ECs, CARs and osteoblasts across four groups of mice. (F) ISG signature scores in Adipo/Osteo-CARs and ECs across four groups of mice. Statistical significances were first determined by Kruskal–Wallis test across four conditions (adjusted P<0.05), followed by post-hoc pairwise tests using Wilcoxon rank sum test. * *P* < 0.05, ** *P* < 0.01, *** *P* < 0.001, **** *P* <0.0001. (G-H) Inferred transcription factor (TF) activities across four groups of mice for adipo-CARs (G) and sinusoidal ECs (H).

Like ECs, CARs of *Perk*^iΔEC^ mice exhibited an enhanced response to IFN (Fig 6D). NK/T cells expressed the highest levels of IFNγ and demonstrated strong communication with adipo-CARs in *Perk*^iΔEC^ mice, but not in DKO mice (supplementary Fig 4E-F). Among stroma subsets, CAR cells had the highest interferon-stimulated gene (ISG) score, calculated from the core enrichment genes of the Hallmark interferon pathway (Fig 6E). The ISG score increased across stroma compartments including adipo-CARs, osteo-CARs, and ECs in *Perk*^iΔEC^ mice but were suppressed in these subsets in DKO mice (Fig 6F). We applied SCENIC gene regulatory network analysis to investigate TF activities from the scRNA-seq data. IFN response TF activity, including *Irf7, Irf1,* and *Stat1,* were all upregulated in *Perk*^iΔEC^ adipo-CARs (Fig 6G), osteo-CARS (not shown), and sinusoidal ECs (Fig 6H). This upregulation was abolished in DKO mice, indicating that these are DLL4-dependent IFN responses promoted by PERK perturbation. In summary, scRNA-seq analysis revealed that the enhanced B lymphocyte recovery in *Perk*^iΔEC^ mice was associated with a strong *Il7-Il7r* signaling axis and heightened IFN response, both of which were markedly attenuated in the absence of DLL4.

### NOTCH3 is the major Notch homolog in CARs and *Notch3^-/-^* mice exhibit impaired post-irradiation lymphoid priming and B progenitor regeneration

CellChat analysis of scRNA-seq data identified *Notch3* as the prime Notch homolog expressed by CARs interacting with DLL4 and Jagged 1 (JAG1) (Fig 7A-B). Importantly, osteo- and adipo-CARs display higher Notch pathway activity than other stroma subsets. Notch activity was enhanced in *Perk*-ablated stroma which had increased DLL4 but was suppressed in DKO stroma (Fig 7C). To determine whether NOTCH3 regulates hematopoiesis, we first examined *Notch3^-/-^* mice under homeostasis and found that steady state hematopoiesis was essentially normal (Fig 7D-F). We then performed reciprocal BM transplantation to determine if NOTCH3 regulates hematopoietic regeneration. While BM cells from *Notch3^-/-^* mice transplanted into WT recipients had normal myeloid and lymphoid lineage development (supplementary Fig 5A-C), *Notch3^-/-^*recipient mice receiving WT BM displayed reduced early B lymphoid recovery and reduced MPP4, CLP, and B progenitor regeneration (Fig 7G-I), suggesting that NOTCH3 in BM stroma may regulate myeloablation-induced B progenitor recovery.

**Figure 7.**
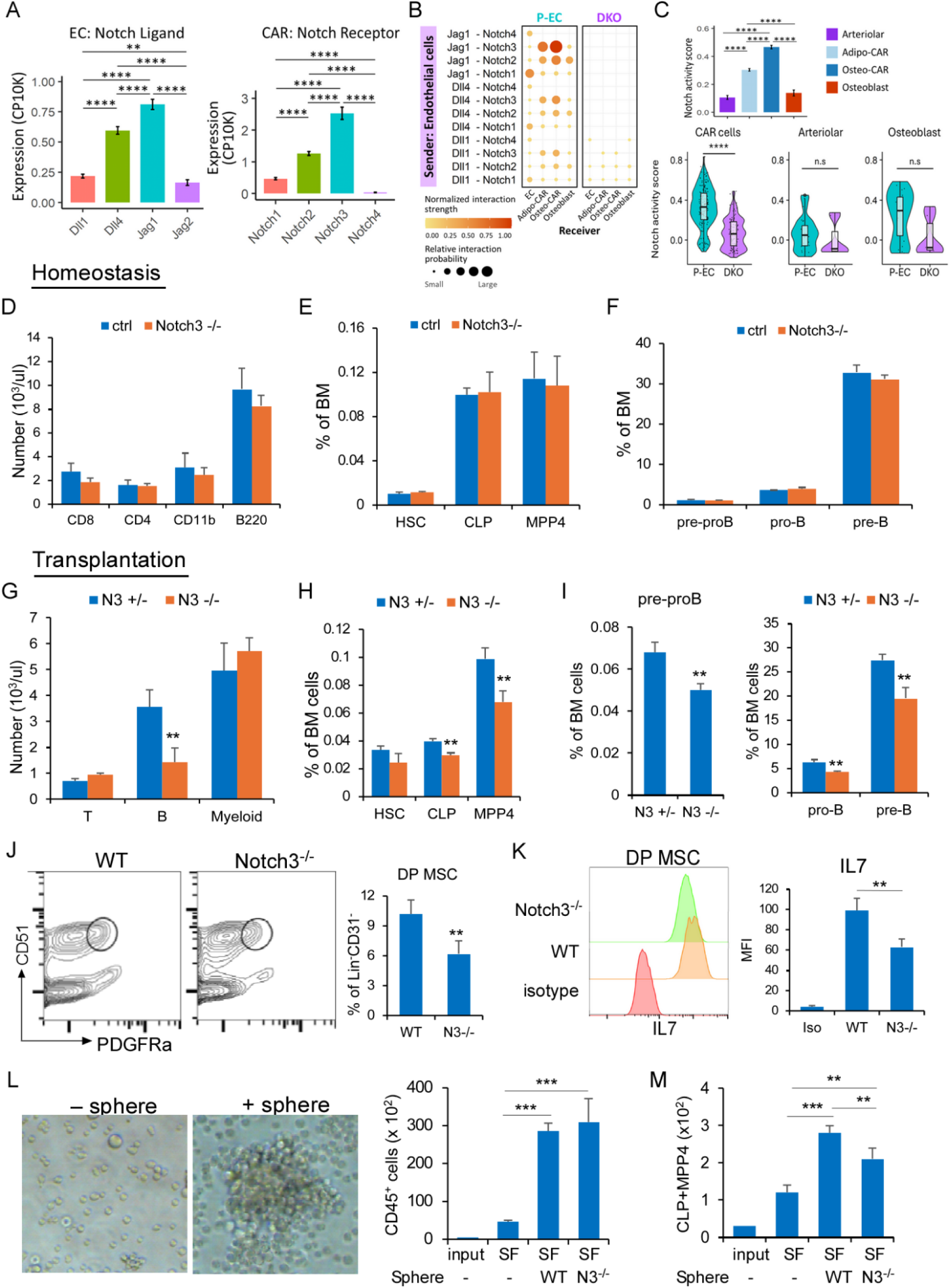
Notch3 deficiency in BM stroma impairs B progenitor regeneration. (A) EC Notch ligand expression and CAR cell Notch receptor expression in the BM. Statistical significances were first determined by Kruskal–Wallis test across four conditions, followed by post-hoc pairwise tests using MAST algorithm. ** *P* < 0.01, **** *P* <0.0001. (B) Dot plot of inferred Notch ligand (sender: EC) - Notch receptor (receiver: EC, CAR-adipo, CAR-osteo, fibroblast/osteoblast) interaction by CellChat analysis. (C) Notch signaling activity score (*Notch3, Notch2, Hes6, Hey1, Hey2, Lfng, Mfng, Maml1* and *Tnc*) in stroma subsets in control mice (top panel) and in stroma subsets of *PERK*^iΔEC^ mice and DKO mice (bottom panel). Statistical significances were first determined by Kruskal–Wallis test across four conditions (adjusted P<0.05), followed by post-hoc pairwise tests using Wilcoxon rank sum test. \*\**P* < 0.01, \*\*\**P* < 0.001, \*\*\*\**P* < 0.0001. (D) Analysis of peripheral blood B cells, T cells, and granulocytes from homeostatic WT and *Notch3^-/-^* mice. Absolute numbers were shown (N=6/group). (E) Frequencies of HSC, CLP and MPP4 (Lin^−^Sca-1^+^c-Kit^+^CD150^-^CD48+Flk2^+^) were determined from WT control and *Notch3^-/-^* mice BM (N=6/group). (F) Numbers of BM B progenitors were determined from WT control and *Notch3^-/-^*mice (n=6/group). (G-I) Analysis of peripheral blood B cells, T cells, and granulocytes at 1 month after lethal irradiation and transplantation of WT donor BM cells in *Notch3^+/-^* and *Notch3^-/-^* recipient mice (G). Frequencies of HSC/CLP/MPP4 (H), pre-proB/preB/proB (I) were determined (n=4/group). Results from D-I are presented as averages ± SD. * *P* < 0.05, ** *P* < 0.01 (J) FACS analysis was caried out on MACSQuant Analyzer 16. Representative FACS profile from 3 similar experiments showing PDGFRα^+^/CD51^+^ (DP) MSCs (Lin^-^Ter119^-^CD31^-^ PDGFRα^+^/CD51^+^) and quantification of DP MSCs in WT or *Notch3^-/-^*mice BM (n=3/group). (K) FACS analysis was caried out on MACSQuant Analyzer 16. Representative FACS profile from 3 similar experiments showing intracellular IL7 expression and quantification of IL7 MFI in DP MSCs isolated from mouse BM eight days after 550 cGy irradiation (n=3/group). (L) A representative photograph showing LK (500/well) cells cultured with SCF and FLT3L (SF) alone or in SF + mesenspheres on day 4 (left) and quantification of expanded total CD45^+^ cells after 5 days of coculture by FACS. (M) Quantification of expanded lymphoid-primed HSPCs (CLP+MPP4) after 5 days of coculture by FACS. Results from J-M are presented as averages ± SD (n=3/group). ** *P* < 0.01, *** *P* < 0.001

To investigate the role of NOTCH3 in the HSC niche, we examined CXCL12 and IL7 production in HSC niche-enriched PDGFRα^+^ and CD51^+^ double-positive (DP) MSCs, which expressed the highest level CXCL12 among all stroma cells (Pinho, Lacombe et al. 2013). There is a slight decrease of DP MSCs in *Notch3^-/-^* mice (Fig 7J), but Notch3-deficiency has no effect on CXCL12 expression (supplemental Fig 5D-E). While WT DP MSCs showed upregulation of IL7 in response to irradiation or IFNβ stimulation (supplemental Fig 5F), Notch3-deficient DP MSCs displayed attenuated post-irradiation IL7 surge but unaltered response to IFNβ (Fig 7K; supplemental Fig 5G). Finally, we assessed MSC niche function using a modified mesensphere coculture assay developed by Frenette’s group (Pinho, Lacombe et al. 2013).

Coculture of WT mesenspheres with BM progenitors (Lin^-^c-kit^+^) in the presence of SCF and FLT3L (SF) led to 6.1-fold and 2.8-fold higher expansion of CD45^+^ cells and lymphoid HSPCs than LK cultured with SF alone. Expansion of lymphoid HSPCs by Notch3-deficient mesenspheres, however, reduced by 25% (Fig 7L-M). Together, these data indicate that NOTCH3 promotes post-irradiation IL7 upregulation and lymphoid priming.

We also investigated the functional significance of JAG1, another major Notch ligand, in post-irradiation hematopoietic regeneration by performing BM transplantation in endothelial *Jag1* deficient mice (*VE-cadherin^ERT2-Cre^/Jag1^F/F^*, or *Jag1^i^*^Δ*EC*^). Unlike *Dll4*^iΔEC^ mice, *Jag1*^iΔEC^ mice displayed no differences in early HSPC recovery (supplemental Fig 6A-B) or long-term HSPC regeneration (not shown) when compared to control mice. Further, there is no significant defect in vascular regeneration in *Jag1*^iΔEC^ mice after irradiation (supplemental Fig 6C). Considered together, we concluded that, in the post-irradiated HSC niche, MSC-expressed NOTCH3 and EC-expressed DLL4 regulates lymphoid priming and B progenitor regeneration.

## Discussion

Enhancing lymphoid recovery is a critical unmet need in HSC transplantation (Savani, Mielke et al. 2007, Ishaqi, Afzal et al. 2008, Le Blanc, Barrett et al. 2009, Yamamoto, Ogusa et al. 2014, Kubo, Imataki et al. 2023). A low absolute lymphocyte count within 3 months of allogeneic HSC transplantation is a major risk factor for worse outcome (Le Blanc, Barrett et al. 2009, Kim, Armand et al. 2015). However, strategies to enhance lymphocyte regeneration are limited, largely due to gaps in our understanding of regenerative mechanisms under myeloablative stress. In this study, we revealed a previously underappreciated role of endothelial ER stress signaling in shaping hematopoietic regeneration. Specifically, activation of endothelial PERK restricts early HSCP regeneration in post-myeloablation transplantation. Our work demonstrated that ablating endothelial *Perk* in irradiated mice increases BM endothelial DLL4 and Notch signaling HSC niche. We revealed that BM stroma NOTCH3, the prime Notch receptor that pairs with EC DLL4, regulates irradiation-induced IL7 upregulation and promotes MSC-supported lymphoid progenitor expansion. Further, single cell analysis reveals that the PERK-DLL4 axis orchestrates the regenerating network by regulating HSC lymphoid commitment, CAR Notch activation, and HSC niche factor expression.

MSCs are essential niche cells for supporting HSC homeostasis (Sugiyama, Kohara et al. 2006, Shen, Tasdogan et al. 2021). Single cell and spatial analysis support a pivotal role for the perivascular niche to instruct lymphoid progenitor development. However, the mechanism that regulates the lymphoid program in the perivascular region in response to stress conditions such as myeloablation has not been elucidated. The close link between the vessels and the perivascular MSC suggests that direct cell-cell interactions are important for HSC niche function. Notch pathway is a fundamental cellular interaction mechanism that controls cell fate decisions via ligand-mediated nuclear translocation of Notch receptor. Several studies including ours have revealed a Notch-regulated mechanism for supporting lymphoid priming in the BM (Yao, Huang et al. 2011, Yu, Saez et al. 2015, Wang, Yu et al. 2017). Here, we provided compelling evidence that endothelial DLL4 and CAR-expressed NOTCH3 are important regulators of lymphoid progenitor regeneration. NOTCH3 is essential for the development of the perivascular MSCs in dental mesenchyme (Koch, Lehal et al. 2013, Pagella, de Vargas Roditi et al. 2021). NOTCH3 also plays a critical role in pathological tissue remodeling (Ramachandran, Dobie et al. 2019, Wei, Korsunsky et al. 2020, Xiang, Pan et al. 2024). In the synovium of rheumatoid arthritis, for example, NOTCH3 drives the perivascular and sublining fibroblast transcriptional gradients initiated from ligand-expressing ECs. Consistently, our work demonstrates that CAR/MSC-expressed NOTCH3 is the prime Notch receptor which engages with endothelial DLL4 to regulate post-transplant lymphoid priming and B progenitor regeneration.

In addition to its Notch-promoting role in CARs, endothelial DLL4 also maintains HSC quiescence and the vessel integrity after transplantation, corroborating our previous work showing increased HSC cycling by blocking DLL4 (Wang, Yu et al. 2015). These roles of DLL4 may explain that HSC and B progenitor recovery were more severely impaired in *Dll4*^iΔEC^ than in *Notch3*-deficient mice. ER stress response is critical for restoration of angiogenic homeostasis after irradiation (Grabham and Sharma 2013). Post-irradiation vasculature in PERK-deficient mice displayed a DLL4-dependent maintenance of organized architecture and vessel integrity, suggesting the interplay of PERK and DLL4 plays a key role in post-irradiation vessel regeneration. Upon PERK activation, phosphorylation of eIF-2α inhibits general translation initiation, which is likely the underlying mechanism of DLL4 de-repression by targeting PERK. Our observation appears to be contradictory to reports showing that DLL4 translation during ER stress can be maintained through PERK-regulated activation of the internal ribosome entry site (IRES) in the 5′-UTR of *DLL4* mRNA (Jaud, Philippe et al. 2019). While exposure to hypoxia incited *DLL4* IRES activity *in vitro*, it is unclear whether translated DLL4 is properly folded and presented on cell surface of ECs, or whether DLL4 degradation is promoted under irradiation-induced ER stress, as protein translation and degradation are tightly regulated by ionizing radiation (Lü, de la Peña et al. 2006). Further studies are needed to illuminate the mechanism by which *Perk* extinction upregulates DLL4 expression under myeloablation. Importantly, endothelial PERK activation and DLL4 suppression are prominent features of BM stroma of leukemia patients undergoing myeloablative treatment. Given the absence of an FDA-approved direct DLL4/NOTCH3-promoting therapeutic strategy, results from our animal studies suggest that pharmacological PERK blockade could improve early lymphoid recovery in post-myeloablation transplantation.

IL7 is essential for B lymphoid lineage commitment and differentiation (Kikuchi, Lai et al. 2005). Increased expression of IL7 in *Perk*^iΔEC^ CARs can drive the accelerated B progenitor recovery. Like the upregulation of IL7 in irradiated thymic stroma and the induction of IL7 by IFN in keratinocytes (Ariizumi, Meng et al. 1995, Toki, Adachi et al. 2003), irradiation and IFNβ increase MSC production of IL7. We found that irradiation-induced increase of IL7 is impaired by *Notch3* deficiency, while *Dll4* ablation in endothelium also led to IL7 reduction based on scRNA-seq analysis. Whether Notch directly regulates IL7 expression remains to be determined. Notably, we identified a strong interaction between T/NK and stroma cells and induction of CAR cell IFN activities by targeting *Perk*. Irf1 is preferentially induced by IFNγ, while Irf1 can induce the expression of ISGs and upregulate type I IFN (Liu, Sanchez et al. 2012).

Increased IFNβ in *Perk*-deleted mice can in turn drives IL7 production in CARs. IFNγ released from T/NK cells is likely regulated by Notch, as T/NK cells show the highest Notch activity among all hematopoietic cells in *Perk*^iΔEC^ BM (not shown) and thus Notch may indirectly regulate CAR IL7 release through IFNγ. Alternatively, IFNγ can be induced by BM cells sensing self-ligands (dsDNA and dsRNA) released upon cell death, as *Perk* extinction increases pyroptosis. IFN plays a context-dependent role in HSCs, with short-term exposure promoting HSC expansion (Essers, Offner et al. 2009, Baldridge, King et al. 2010, Demerdash, Kain et al. 2021). Together, these findings suggest that *Perk* ablation elicits DLL4-dependent coordinated IFN response, Notch activation, and IL7 release from CARs, which synergistically promote HSPC and B progenitor regeneration.

In summary, we identify that vascular PERK-DLL4 axis and CAR-expressing NOTCH3 as central regulators of a specialized “pre-B niche” that governs lymphoid regeneration following transplantation. By linking endothelial ER stress, vascular–stromal Notch signaling, IFN response, and IL7 production, our findings not only uncover fundamental mechanisms of lymphoid regeneration but also highlight PERK inhibition as a promising therapeutic strategy to improve immune recovery after myeloablative transplantation.

## Materials and Methods

### Mice and treatment

Animal studies were approved by the IACUC. B6;129S1-Notch3tm1Grid/J mice (Krebs, Xue et al. 2003) (JAX^®^#010547) were backcrossed to C57BL/6 mice and confirmed to be 100% C57BL/6 background by genome scan analysis. Previously described floxed mice (Perk^F/F^ (Zhang, McGrath et al. 2002), Jag1^F/F^ (Basu, Barbur et al. 2018), Dll4^F/F^) and VE-cadherin^ERT2-Cre^ mice (Sörensen, Adams et al. 2009) were maintained on the C57BL6 background and crossed to generate inducible endothelial-specific knockout of *Perk*, *Dll4*, or *Jag1* by five consecutive doses of tamoxifen. PERK and DLL4 expression were assessed by anti-PERK (BS-2469R; Bioss Antibodies Inc.) and anti-DLL4 (HMD4-1; Thermo Fisher) through FACS. For pharmacologic treatments, mice received GSK2656157 (Selleckchem) (50 mg/kg in 10% DMSO and 90% corn oil) through oral gavage twice daily starting on day 3 after transplantation for 12 days. BI09 (Tang, Ranatunga et al. 2014) was administered daily by intraperitoneal injection (25 mg/kg) for 5 days, starting one day before irradiation.

### BM transplantation and analysis of peripheral and bone marrow hematopoiesis

BM transplantation was performed as described previously (Wang, Zimmerman et al. 2016). Briefly, 2.0 x10^6^ total BM cells from WT donor mice (Ly5.1) were transferred retro-orbitally to lethally irradiated (950 cGy Cesium-137 irradiator or 850 cGy X-ray irradiator; lethal for at least 50% of mice by day 30 (LD50/30) in our facility) WT recipients (Ly5.2). For competitive secondary and tertiary transplantation, BM cells from primary or secondary transplant recipients (Ly5.1) were assessed for HSC frequency, adjusted for the equivalent HSC numbers within 1.0 x10^6^ competitive bone marrow cells (Ly5.2), and infused together with 1.0 x10^6^ competitive cells into lethally irradiated secondary or tertiary recipients (Ly5.2). Non-lethal irradiation (550 cGy X-ray) was performed in assessing endothelial DLL4 expression, whole mount and intravital imaging, and scRNA-sequencing. Chemotherapy with 5-FU was performed by a single IP injection of 5-FU (150 mg/kg). BM reconstitution was analyzed at d21 and monthly thereafter by manual peripheral blood white blood cell counts (hemocytometer) and FACS immunophenotyping. Three independent groups of transplanted mice were analyzed separately, and results were pooled.

For BM HSPC reconstitution analysis, BM nucleated cells collected from two femurs and two tibias were stained with a cocktail of biotinylated lineage antibodies followed by analysis of HSPC (Lin^-^Sca1^+^c-kit^+^). Antibodies were purchased from BD (San Jose, CA), eBioscience (San Diego, CA), and Biolegend (San Diego, CA) and included those against the following antigens: CD4 (RM4-5), CD8α (53-6.7), B220 (RA3-6B2), CD11b (M1/70), Gr-1 (RB6-8C5), TER119 (TER-119), c-KIT (2B8), Sca1 (D7), CD150 (9D1), CD34 (RAM34), CD16/CD32 (2.4G2), and Flt3 (A2F10). Analysis of HSPC (Lin^-^Sca1^+^c-kit^+^) was achieved by gating on lineage^-^ (CD4^-^CD8^-^ B220^-^CD11b^-^Gr-1^-^TER119^-^) cells. Analysis of CAR and endothelial cells was achieved by gating on CD45^-^TER119^-^CD71^-^ cells. Antibodies used for analysis are CD31 (390), CD71 (RI7217), PERK (C33E10), p-PERK (T980) (16F8), DLL4 (HMD4-1), Cxcl12 (79018), and IL7 (PA5-79509). FACS analyses were performed on CytoFLEX (Case Western Reserve University), BD LSRII or MACSQuant Analyzer 16 (Houston Methodist Hospital) after washing cells with PBS. For p-EPRK and Cxcl12 analysis, BM cells were incubated with CD31, TER119, and CD45 for 20 min on ice and then fixed (BD Biosciences #557870) and permeabilized (BD Biosciences #554723) for 20 min at room temperature. Cells were then incubated with the p-PERK or Cxcl12 antibodies for 20-30 min. For IL7 analysis, cells were incubated with anti-IL7 antibody (1:100, Invitrogen #PA5-79509) at room temperature for 45 min after fixation/permeabilization followed by staining with FITC conjugated anti-rabbit antibody (Biolegend #406403) at room temperature for 20 min.

### MSC culture and analysis

BM stroma cells were collected and enriched as described (Pinho, Lacombe et al. 2013). Total BM cells were flushed from two femurs and two tibias, resuspended in MSC medium (MesenCult™ Expansion Kit, Catalog # 05513, StemCell) after red cell lysis, and plated in 6-well plates and allowed to adhere. Non-adherent hematopoietic cells were removed daily for the first 3 days and then twice per week with MSC medium. MSCs were harvested for FACS analysis after reaching 75% confluence or further enriched by sorting. Mesensphere formation was carried out using a modified protocol (Mendez-Ferrer, Michurina et al. 2010). Briefly, MSCs cultured in 2D were digested with 0.05% trypsin, neutralized with 10 volumes of 10% FBS and resuspended in MSC 3D culture medium at clonal density in ultra-low attachment 96-well plates (Corning #3474). The MSC 3D culture medium contained 15% chicken embryo extract, 0.1 mM β-mercaptoethanol, 1% non-essential amino acids (Sigma), 1% N2, 2% B27 supplements (Gibco), and 20 ng/mL each of fibroblast growth factor (FGF)-basic, insulin-like growth factor-1 (IGF-1), epidermal growth factor (EGF), platelet-derived growth factor (PDGF) and oncostatin M (OSM) (Peprotech) in DMEM/F12 (1:1) / human endothelial (Gibco) (1:2). MSC cultures were maintained at 37 °C with 5% CO_2_ in a water-jacketed incubator and left untouched for 1 week to allow mesensphere formation, with half medium changes performed weekly. For coculture assays, mouse BM Lin^-^c-kit^+^ (LK) cells were prepared using CD117 microbeads following anti– biotin magnetic microbeads (Miltenyi Biotec) after incubation with a cocktail of biotinylated antibodies (Gr-1, CD11b, CD4, CD8, NK1.1, B220, and TER119). Approximately 500 Lin^-^c-kit^+^ cells were cocultured with ∼20 mesenspheres in medium (StemSpan; STEMCELL Technologies) supplemented with 50 ng/mL of SCF and 100 ng/mL of Flt3L (R&D Systems) for 5 days at 37°C. For analysis of IL7 in MSCs, cells were stimulated with or without IFNβ (1500 IU/ml, 4 h, 37 °C) followed by incubation with anti-IL7 antibody as described above.

### Immunohistochemistry

Studies involving human marrow specimens were approved by the Institutional Research Board (IRB) of the Houston Methodist Research Institute. Expression of p-PERK and DLL4 was evaluated by immunohistochemistry (IHC). Tissues sections were incubated overnight at 4°C with the primary antibodies against p-PERK (1:200, Cell Signaling) DLL4 (21584-1-AP, 1:400, Proteintech). Slides were scanned using the MoticEasyScan Pro 6 (Motic) and acquired images were analyzed with the accompanying software.

### scRNA-seq library construction

For scRNA-seq, mice were irradiated with a sublethal 550 cGy (RS 2000) sufficient to induce evident damage to the BM vasculature and myelosuppression (Himburg, Sasine et al. 2016). Single-cell RNA sequencing libraries were prepared according to manufacturer’s instructions (10X Genomics, 3’ V3) using pooled mouse marrow cells (n=4-5/genotype; female and male mixed) collected on day 11-12, a time point when the collapsed vascular network has recovered by whole mount immunostaining and leakiness assay in the wild type mice based on other reports and our own analysis (Himburg, Sasine et al. 2016). The scRNA-seq libraries were generated by Chromium nest GEM Single Cell 3’ v3.1 kit (PN-1000263, 10x Genomics, Pleasanton, CA) according to the manufacturer’s protocol. Pre-made libraries were sequenced on a Novaseq 6000 (Illumina) instrument to obtain a minimum of 30,000 paired end reads per cell.

### scRNA-seq data quality control and cell annotation

The FASTQ data were processed using CellRanger (version 7.1.0), and downstream analyses were conducted in R with Seurat package (version 5.0.1) (Hao, Stuart et al. 2024). Specifically, doublets were eliminated using DoubletFinder (McGinnis, Murrow et al. 2019), and ambient RNA counts were corrected with DecontX (Yang, Corbett et al. 2020). Low-quality cells, characterized by high mitochondrial content and low gene detection rates, were also filtered out. The quality control workflow resulted in 38,031 total cells, with each condition ranging from 8,062 to 10,366 cells. Gene counts were SCTransformed and identified top 3,000 highly variable genes (HVGs) to obtain principal components (PCs). All libraries were integrated by Harmony using top 30 PCs and then clustered by the Louvain method (Korsunsky, Millard et al. 2019). Cell clusters were annotated as fourteen cell population based on known cellular markers. SingleR and reference mapping approaches were used to verify cell type annotations (using bone marrow single cell atlas) to ensure accurate annotation of HSPCs/cycling progenitors under irradiation conditions. Stromal cells were further subset (n=2798) and re-clustered following the above-mentioned approach. The refined clustering identifies five stromal cell subtypes. Parameter titrations (data not shown) were applied using different numbers of HVGs, PCs, and clustering resolutions to ensure a robust phenotyping of stromal cell types.

### scRNA-seq computational analysis

The integrated scRNA-seq data were used to identify differentially expressed genes (DEGs) using MAST algorithm (Finak, McDavid et al. 2015), followed by pathway analysis. MAST is a two-part, generalized linear modeling approach that can robustly detect DEGs and calculate fold changes of gene expression, even with varying numbers of single cells in different groups and for lowly expressed genes. MAST result reported log2 fold-change (L2FC) and Benjamini-Hochberg (BH) adjusted p-value (*P*_adj_). For gene expression bar plots, gene expression data were first normalized to a unit of count per 10,000 (CP10K) and grouped by experimental conditions. Group-wise statistical comparisons were reported using the results from MAST algorithm. Then, all DEGs were ranked based on L2FC x Truncated(-LOG10(P_adj_)) for Gene Set Enrichment Analysis (GSEA). Gene Ontology (GO) database was used for functional annotation in the ClusterProfiler package (Wu, Hu et al. 2021). Normalized Enrichment Score, *P* and *P*_adj_ values were used for identifying significantly up or down regulated pathways. Cell-cell communication was analyzed using CellChat based on log-normalized counts (Jin, Plikus et al. 2024). CellChat inferred cell-type-wise communication probability for each experimental group and ligand-receptor (LR) pairs. The bubble and circle plots visualized the cell-cell communication strength for selected signaling pathway and LR pairs. Gene regulatory networks were analyzed using pySCENIC (Van de Sande, Flerin et al. 2020), which computationally inferred the activities of transcription factors (TFs) at single cell resolution from gene expression data. Wilcoxon rank sum test was deployed to statistically compare the differentially activated TFs across experimental conditions.

### Multiphoton imaging and analysis for vessel leakiness

Intravital 2-photon imaging preparation was performed as previously described (Myers, Huang et al. 2010, Wang, Zimmerman et al. 2016). Mice were anaesthetized with isoflurane and placed in a stereotactic holder. The skin above the calvarium was carefully removed and a well was created with dental acrylic around the calvarium to hold sterile saline for imaging. The mouse was positioned under the microscope to image the frontal bone near the junction of the interfrontal and coronal sutures. Vessel dye containing 1 mg each of 2000 kD FITC-Dextran or 70 kD TRITC-Dextran (Sigma-Aldrich) was injected retro-orbitally immediately before imaging.

Images were acquired on a Leica SP5 microscope equipped with a Coherent Chameleon Ti/Sapphire laser tuned to 800 nm and a 4-channel-NDD detector while mice were under inhaled anesthesia (1%–2% isoflurane) on a warmed microscope stage (37^0^C). Three dimensional images consisting of XY: 775 mm x 775 mm and Z: 140 mm (5 mm/step), were obtained every 30 seconds for 30 minutes. Images were analyzed with Imaris software (Oxford Instruments). Datasets were drift-corrected, smoothed, and surfaces were generated on both FITC and TRITC channels. The corresponding total fluorescent volumes for each tracer were calculated at each timepoint and normalized to the first timepoint.

### Whole-mount immunostaining

To analyze the vascular network, 10 μg each of Alexa Fluor 647 anti-mouse CD31 and Alexa Fluor 647 anti-mouse CD144 (BioLegend) were administered intravenously 10 min before euthanasia, as described (Kunisaki, Bruns et al. 2013, Lucas, Scheiermann et al. 2013). Sternal bone fragments were fixed in ice cold 4% PFA for 3h. Whole-mount fragments were directly imaged on a Leica SP5 inverted confocal imaging system. Fluorescent emissions were collected using internal detectors set to 494 – 538 nm (GFP), 553 – 617 nm (PE), and 644 – 710 nm (AF647). High resolution three-dimensional (xyz) scans were performed using a Leica 10x objective (N.A. 0.4), with XYZ voxel sizes of 1.5 μm x 1.5 μm x 5 μm. Images were analyzed using Imaris software (Bitplane, Inc, Belfast, UK). Volume data for each image was determined by generating surfaces in Imaris based on the corresponding fluorescent signals. The overall volume of the image was used to normalize the data and to determine percentages. **Statistical analysis**

Statistics of all paired experiments were analyzed by two-tailed Student’s *t* test or two-way ANOVA. Statistics of scRNA-seq were described in the figure legends.

## Acknowledgement

This study was supported in part by research funding from HL103827 and CA222064 to LZ.

## References

Ariizumi, K., Y. Meng, P. R. Bergstresser and A. Takashima (1995). “IFN-gamma-dependent IL-7 gene regulation in keratinocytes.” J Immunol 154(11): 6031–6039.

Baccin, C., J. Al-Sabah, L. Velten, P. M. Helbling, F. Grünschläger, P. Hernández-Malmierca, C. Nombela-Arrieta, L. M. Steinmetz, A. Trumpp and S. Haas (2020). “Combined single-cell and spatial transcriptomics reveal the molecular, cellular and spatial bone marrow niche organization.” Nat Cell Biol 22(1): 38–48.

Baldridge, M. T., K. Y. King, N. C. Boles, D. C. Weksberg and M. A. Goodell (2010). “Quiescent haematopoietic stem cells are activated by IFN-gamma in response to chronic infection.” Nature 465(7299): 793–797.

Basu, S., I. Barbur, A. Calderon, S. Banerjee and A. Proweller (2018). “Notch signaling regulates arterial vasoreactivity through opposing functions of Jagged1 and Dll4 in the vessel wall.” Am J Physiol Heart Circ Physiol 315(6): H1835–h1850.

Benedito, R., C. Roca, I. Sorensen, S. Adams, A. Gossler, M. Fruttiger and R. H. Adams (2009). “The notch ligands Dll4 and Jagged1 have opposing effects on angiogenesis.” Cell 137(6): 1124–1135.

Butler, J. M., D. J. Nolan, E. L. Vertes, B. Varnum-Finney, H. Kobayashi, A. T. Hooper, M. Seandel, K. Shido, I. A. White, M. Kobayashi, L. Witte, C. May, C. Shawber, Y. Kimura, J. Kitajewski, Z. Rosenwaks, I. D. Bernstein and S. Rafii (2010). “Endothelial cells are essential for the self-renewal and repopulation of Notch-dependent hematopoietic stem cells.” Cell Stem Cell 6(3): 251–264.

Calvi, L. M. and D. C. Link (2015). “The hematopoietic stem cell niche in homeostasis and disease.” Blood 126(22): 2443–2451.

Comazzetto, S., M. M. Murphy, S. Berto, E. Jeffery, Z. Zhao and S. J. Morrison (2019). “Restricted Hematopoietic Progenitors and Erythropoiesis Require SCF from Leptin Receptor+ Niche Cells in the Bone Marrow.” Cell Stem Cell 24(3): 477–486.e476.

Cordeiro Gomes, A., T. Hara, V. Y. Lim, D. Herndler-Brandstetter, E. Nevius, T. Sugiyama, S. Tani-Ichi, S. Schlenner, E. Richie, H. R. Rodewald, R. A. Flavell, T. Nagasawa, K. Ikuta and J. P. Pereira (2016). “Hematopoietic Stem Cell Niches Produce Lineage-Instructive Signals to Control Multipotent Progenitor Differentiation.” Immunity 45(6): 1219–1231.

Demerdash, Y., B. Kain, M. A. G. Essers and K. Y. King (2021). “Yin and Yang: The dual effects of interferons on hematopoiesis.” Exp Hematol 96: 1–12.

Ding, L. and S. J. Morrison (2013). “Haematopoietic stem cells and early lymphoid progenitors occupy distinct bone marrow niches.” Nature 495(7440): 231–235.

Ding, L., T. L. Saunders, G. Enikolopov and S. J. Morrison (2012). “Endothelial and perivascular cells maintain haematopoietic stem cells.” Nature 481(7382): 457–462.

Ding, W., L. Yang, M. Zhang and Y. Gu (2012). “Reactive oxygen species-mediated endoplasmic reticulum stress contributes to aldosterone-induced apoptosis in tubular epithelial cells.” Biochem Biophys Res Commun 418(3): 451–456.

Essers, M. A., S. Offner, W. E. Blanco-Bose, Z. Waibler, U. Kalinke, M. A. Duchosal and A. Trumpp (2009). “IFNalpha activates dormant haematopoietic stem cells in vivo.” Nature 458(7240): 904–908.

Finak, G., A. McDavid, M. Yajima, J. Deng, V. Gersuk, A. K. Shalek, C. K. Slichter, H. W. Miller, M. J. McElrath, M. Prlic, P. S. Linsley and R. Gottardo (2015). “MAST: a flexible statistical framework for assessing transcriptional changes and characterizing heterogeneity in single-cell RNA sequencing data.” Genome Biology 16(1): 278.

Grabham, P. and P. Sharma (2013). “The effects of radiation on angiogenesis.” Vasc Cell 5(1): 19.

Greenbaum, A., Y. M. Hsu, R. B. Day, L. G. Schuettpelz, M. J. Christopher, J. N. Borgerding, T. Nagasawa and D. C. Link (2013). “CXCL12 in early mesenchymal progenitors is required for haematopoietic stem-cell maintenance.” Nature 495(7440): 227–230.

Greenberg, Z. J., L. C. Paracatu, D. A. Monlish, Q. Dong, M. Rettig, N. Roundy, R. Gaballa, W. Li, W. Yang, C. J. Luke and L. G. Schuettpelz (2023). “The tetraspanin CD53 protects stressed hematopoietic stem cells via promotion of DREAM complex-mediated quiescence.” Blood 141(10): 1180–1193.

Hao, Y., T. Stuart, M. H. Kowalski, S. Choudhary, P. Hoffman, A. Hartman, A. Srivastava, G. Molla, S. Madad, C. Fernandez-Granda and R. Satija (2024). “Dictionary learning for integrative, multimodal and scalable single-cell analysis.” Nature Biotechnology 42(2): 293–304.

Himburg, H. A., J. Sasine, X. Yan, J. Kan, H. Dressman and J. P. Chute (2016). “A Molecular Profile of the Endothelial Cell Response to Ionizing Radiation.” Radiat Res 186(2): 141–152.

Ishaqi, M. K., S. Afzal, A. Dupuis, J. Doyle and A. Gassas (2008). “Early lymphocyte recovery post-allogeneic hematopoietic stem cell transplantation is associated with significant graft-versus-leukemia effect without increase in graft-versus-host disease in pediatric acute lymphoblastic leukemia.” Bone Marrow Transplant 41(3): 245–252.

Jaud, M., C. Philippe, L. Van Den Berghe, C. Ségura, L. Mazzolini, S. Pyronnet, H. Laurell and C. Touriol (2019). “The PERK Branch of the Unfolded Protein Response Promotes DLL4 Expression by Activating an Alternative Translation Mechanism.” Cancers (Basel) 11(2).

Jin, S., M. V. Plikus and Q. Nie (2024). “CellChat for systematic analysis of cell–cell communication from single-cell transcriptomics.” Nature Protocols.

Kikuchi, K., A. Y. Lai, C. L. Hsu and M. Kondo (2005). “IL-7 receptor signaling is necessary for stage transition in adult B cell development through up-regulation of EBF.” J Exp Med 201(8): 1197–1203.

Kim, D. H., J. G. Kim, S. K. Sohn, W. J. Sung, J. S. Suh, K. S. Lee and K. B. Lee (2004). “Clinical impact of early absolute lymphocyte count after allogeneic stem cell transplantation.” Br J Haematol 125(2): 217–224.

Kim, H. T., P. Armand, D. Frederick, E. Andler, C. Cutler, J. Koreth, E. P. Alyea, 3rd, J. H. Antin, R. J. Soiffer, J. Ritz and V. T. Ho (2015). “Absolute lymphocyte count recovery after allogeneic hematopoietic stem cell transplantation predicts clinical outcome.” Biol Blood Marrow Transplant 21(5): 873–880.

Koch, U., R. Lehal and F. Radtke (2013). “Stem cells living with a Notch.” Development 140(4): 689–704.

Korsunsky, I., N. Millard, J. Fan, K. Slowikowski, F. Zhang, K. Wei, Y. Baglaenko, M. Brenner, P.-r. Loh and S. Raychaudhuri (2019). “Fast, sensitive and accurate integration of single-cell data with Harmony.” Nature Methods 16(12): 1289–1296.

Krebs, L. T., Y. Xue, C. R. Norton, J. P. Sundberg, P. Beatus, U. Lendahl, A. Joutel and T. Gridley (2003). “Characterization of Notch3-deficient mice: normal embryonic development and absence of genetic interactions with a Notch1 mutation.” Genesis 37(3): 139–143.

Kubo, H., O. Imataki, T. Fukumoto, T. Ishida, Y. H. Kubo, J. I. Kida, M. Uemura, H. Fujita and N. Kadowaki (2023). “Prognostic impact of the simple L-index and absolute lymphocyte count early after allogeneic hematopoietic stem cell transplantation.” Cytotherapy 25(4): 415–422.

Kunisaki, Y., I. Bruns, C. Scheiermann, J. Ahmed, S. Pinho, D. Zhang, T. Mizoguchi, Q. Wei, D. Lucas, K. Ito, J. C. Mar, A. Bergman and P. S. Frenette (2013). “Arteriolar niches maintain haematopoietic stem cell quiescence.” Nature 502(7473): 637–643.

Le Blanc, K., A. J. Barrett, M. Schaffer, H. Hägglund, P. Ljungman, O. Ringdén and M. Remberger (2009). “Lymphocyte recovery is a major determinant of outcome after matched unrelated myeloablative transplantation for myelogenous malignancies.” Biol Blood Marrow Transplant 15(9): 1108–1115.

Liu, C., Q. Chen, Y. Shang, L. Chen, J. Myers, A. Awadallah, J. Sun, S. Yu, K. Umphred-Wilson, D. Che, Y. Dou, L. Li, P. Wearsch, D. Ramírez-Bergeron, R. Beck, W. Xin, G. Jin, S. Adoro and L. Zhou (2022). “Endothelial PERK-ATF4-JAG1 axis activated by T-ALL remodels bone marrow vascular niche.” Theranostics 12(6): 2894–2907.

Liu, S. Y., D. J. Sanchez, R. Aliyari, S. Lu and G. Cheng (2012). “Systematic identification of type I and type II interferon-induced antiviral factors.” Proc Natl Acad Sci U S A 109(11): 4239–4244.

Lü, X., L. de la Peña, C. Barker, K. Camphausen and P. J. Tofilon (2006). “Radiation-induced changes in gene expression involve recruitment of existing messenger RNAs to and away from polysomes.” Cancer Res 66(2): 1052–1061.

Lucas, D., C. Scheiermann, A. Chow, Y. Kunisaki, I. Bruns, C. Barrick, L. Tessarollo and P. S. Frenette (2013). “Chemotherapy-induced bone marrow nerve injury impairs hematopoietic regeneration.” Nat Med 19(6): 695–703.

Malhotra, J. D. and R. J. Kaufman (2007). “Endoplasmic reticulum stress and oxidative stress: a vicious cycle or a double-edged sword?” Antioxid Redox Signal 9(12): 2277–2293.

McGinnis, C. S., L. M. Murrow and Z. J. Gartner (2019). “DoubletFinder: Doublet Detection in Single-Cell RNA Sequencing Data Using Artificial Nearest Neighbors.” Cell Systems 8(4): 329–337.e324.

Mendez-Ferrer, S., T. V. Michurina, F. Ferraro, A. R. Mazloom, B. D. Macarthur, S. A. Lira, D. T. Scadden, A. Ma’ayan, G. N. Enikolopov and P. S. Frenette (2010). “Mesenchymal and haematopoietic stem cells form a unique bone marrow niche.” Nature 466(7308): 829–834.

Mikkelsen, R. B. and P. Wardman (2003). “Biological chemistry of reactive oxygen and nitrogen and radiation-induced signal transduction mechanisms.” Oncogene 22(37): 5734–5754.

Morrison, S. J. and D. T. Scadden (2014). “The bone marrow niche for haematopoietic stem cells.” Nature 505(7483): 327–334.

Myers, J., Y. Huang, L. Wei, Q. Yan, A. Huang and L. Zhou (2010). “Fucose-deficient hematopoietic stem cells have decreased self-renewal and aberrant marrow niche occupancy.” Transfusion 50(12): 2660–2669.

Pagella, P., L. de Vargas Roditi, B. Stadlinger, A. E. Moor and T. A. Mitsiadis (2021). “Notch signaling in the dynamics of perivascular stem cells and their niches.” Stem Cells Transl Med 10(10): 1433–1445.

Patel, B., Y. Zhou, R. L. Babcock, F. Ma, M. A. Zal, D. Kumar, Y. B. Medik, L. M. Kahn, J. E. Pineda, E. M. Park, S. M. Schneider, X. Tang, M. G. Raso, C. R. Jeter, T. Zal, K. Clise-Dwyer, K. Keyomarsi, F. G. Giancotti, S. Colla and S. S. Watowich (2024). “STAT3 protects hematopoietic stem cells by preventing activation of a deleterious autocrine type-I interferon response.” Leukemia 38(5): 1143–1155.

Pinho, S. and P. S. Frenette (2019). “Haematopoietic stem cell activity and interactions with the niche.” Nat Rev Mol Cell Biol 20(5): 303–320.

Pinho, S., J. Lacombe, M. Hanoun, T. Mizoguchi, I. Bruns, Y. Kunisaki and P. S. Frenette (2013). “PDGFRα and CD51 mark human nestin+ sphere-forming mesenchymal stem cells capable of hematopoietic progenitor cell expansion.” J Exp Med 210(7): 1351–1367.

Ramachandran, P., R. Dobie, J. R. Wilson-Kanamori, E. F. Dora, B. E. P. Henderson, N. T. Luu, J. R. Portman, K. P. Matchett, M. Brice, J. A. Marwick, R. S. Taylor, M. Efremova, R. Vento-Tormo, N. O. Carragher, T. J. Kendall, J. A. Fallowfield, E. M. Harrison, D. J. Mole, S. J. Wigmore, P. N. Newsome, C. J. Weston, J. P. Iredale, F. Tacke, J. W. Pollard, C. P. Ponting, J. C. Marioni, S. A. Teichmann and N. C. Henderson (2019). “Resolving the fibrotic niche of human liver cirrhosis at single-cell level.” Nature 575(7783): 512–518.

Ranatunga, S., C. H. Tang, C. W. Kang, C. L. Kriss, B. J. Kloppenburg, C. C. Hu and J. R. Del Valle (2014). “Synthesis of novel tricyclic chromenone-based inhibitors of IRE-1 RNase activity.” J Med Chem 57(10): 4289–4301.

Sato, A., N. Kamio, A. Yokota, Y. Hayashi, A. Tamura, Y. Miura, T. Maekawa and H. Hirai (2020). “C/EBPβ isoforms sequentially regulate regenerating mouse hematopoietic stem/progenitor cells.” Blood Adv 4(14): 3343–3356.

Savani, B. N., S. Mielke, K. Rezvani, A. Montero, A. S. Yong, L. Wish, J. Superata, R. Kurlander, A. Singh, R. Childs and A. J. Barrett (2007). “Absolute lymphocyte count on day 30 is a surrogate for robust hematopoietic recovery and strongly predicts outcome after T cell-depleted allogeneic stem cell transplantation.” Biol Blood Marrow Transplant 13(10): 1216–1223.

Shen, B., A. Tasdogan, J. M. Ubellacker, J. Zhang, E. D. Nosyreva, L. Du, M. M. Murphy, S. Hu, Y. Yi, N. Kara, X. Liu, S. Guela, Y. Jia, V. Ramesh, C. Embree, E. C. Mitchell, Y. C. Zhao, L. A. Ju, Z. Hu, G. M. Crane, Z. Zhao, R. Syeda and S. J. Morrison (2021). “A mechanosensitive peri-arteriolar niche for osteogenesis and lymphopoiesis.” Nature 591(7850): 438–444.

Silberstein, L., K. A. Goncalves, P. V. Kharchenko, R. Turcotte, Y. Kfoury, F. Mercier, N. Baryawno, N. Severe, J. Bachand, J. A. Spencer, A. Papazian, D. Lee, B. R. Chitteti, E. F. Srour, J. Hoggatt, T. Tate, C. Lo Celso, N. Ono, S. Nutt, J. Heino, K. Sipilä, T. Shioda, M. Osawa, C. P. Lin, G. F. Hu and D. T. Scadden (2016). “Proximity-Based Differential Single-Cell Analysis of the Niche to Identify Stem/Progenitor Cell Regulators.” Cell Stem Cell 19(4): 530–543.

Solaimani Kartalaei, P., T. Yamada-Inagawa, C. S. Vink, E. de Pater, R. van der Linden, J. Marks-Bluth, A. van der Sloot, M. van den Hout, T. Yokomizo, M. L. van Schaick-Solernó, R. Delwel, J. E. Pimanda, I. W. F. van and E. Dzierzak (2015). “Whole-transcriptome analysis of endothelial to hematopoietic stem cell transition reveals a requirement for Gpr56 in HSC generation.” J Exp Med 212(1): 93–106.

Sörensen, I., R. H. Adams and A. Gossler (2009). “DLL1-mediated Notch activation regulates endothelial identity in mouse fetal arteries.” Blood 113(22): 5680–5688.

Spangrude, G. J., S. Heimfeld and I. L. Weissman (1988). “Purification and characterization of mouse hematopoietic stem cells.” Science 241(4861): 58–62.

Sugiyama, T., H. Kohara, M. Noda and T. Nagasawa (2006). “Maintenance of the hematopoietic stem cell pool by CXCL12-CXCR4 chemokine signaling in bone marrow stromal cell niches.” Immunity 25(6): 977–988.

Tang, C. H., S. Ranatunga, C. L. Kriss, C. L. Cubitt, J. Tao, J. A. Pinilla-Ibarz, J. R. Del Valle and C. C. Hu (2014). “Inhibition of ER stress-associated IRE-1/XBP-1 pathway reduces leukemic cell survival.” J Clin Invest 124(6): 2585–2598.

Termini, C. M., A. Pang, T. Fang, M. Roos, V. Y. Chang, Y. Zhang, N. J. Setiawan, L. Signaevskaia, M. Li, M. M. Kim, O. Tabibi, P. K. Lin, J. P. Sasine, A. Chatterjee, R. Murali, H. A. Himburg and J. P. Chute (2021). “Neuropilin 1 regulates bone marrow vascular regeneration and hematopoietic reconstitution.” Nat Commun 12(1): 6990.

Tikhonova, A. N., I. Dolgalev, H. Hu, K. K. Sivaraj, E. Hoxha, A. Cuesta-Dominguez, S. Pinho, I. Akhmetzyanova, J. Gao, M. Witkowski, M. Guillamot, M. C. Gutkin, Y. Zhang, C. Marier, C. Diefenbach, S. Kousteni, A. Heguy, H. Zhong, D. R. Fooksman, J. M. Butler, A. Economides, P. S. Frenette, R. H. Adams, R. Satija, A. Tsirigos and I. Aifantis (2019). “The bone marrow microenvironment at single-cell resolution.” Nature 569(7755): 222–228.

Toki, J., Y. Adachi, T. Jin, T. Fan, K. Takase, Z. Lian, H. Hayashi, M. E. Gershwin and S. Ikehara (2003). “Enhancement of IL-7 following irradiation of fetal thymus.” Immunobiology 207(4): 247–258.

Van de Sande, B., C. Flerin, K. Davie, M. De Waegeneer, G. Hulselmans, S. Aibar, R. Seurinck, W. Saelens, R. Cannoodt, Q. Rouchon, T. Verbeiren, D. De Maeyer, J. Reumers, Y. Saeys and S. Aerts (2020). “A scalable SCENIC workflow for single-cell gene regulatory network analysis.” Nature Protocols 15(7): 2247–2276.

van Galen, P., A. Kreso, N. Mbong, D. G. Kent, T. Fitzmaurice, J. E. Chambers, S. Xie, E. Laurenti, K. Hermans, K. Eppert, S. J. Marciniak, J. C. Goodall, A. R. Green, B. G. Wouters, E. Wienholds and J. E. Dick (2014). “The unfolded protein response governs integrity of the haematopoietic stem-cell pool during stress.” Nature 510(7504): 268–272.

Voit, R. A., L. Tao, F. Yu, L. D. Cato, B. Cohen, T. J. Fleming, M. Antoszewski, X. Liao, C. Fiorini, S. K. Nandakumar, L. Wahlster, K. Teichert, A. Regev and V. G. Sankaran (2023). “A genetic disorder reveals a hematopoietic stem cell regulatory network co-opted in leukemia.” Nat Immunol 24(1): 69–83.

Wang, W., S. Yu, J. Myers, Y. Wang, W. W. Xin, M. Albakri, A. W. Xin, M. Li, A. Y. Huang, W. Xin, C. W. Siebel, H. M. Lazarus and L. Zhou (2017). “Notch2 blockade enhances hematopoietic stem cell mobilization and homing.” Haematologica 102(10): 1785–1795.

Wang, W., S. Yu, G. Zimmerman, Y. Wang, J. Myers, V. W. Yu, D. Huang, X. Huang, J. Shim, Y. Huang, W. Xin, P. Qiao, M. Yan, W. Xin, D. T. Scadden, P. Stanley, J. B. Lowe, A. Y. Huang, C. W. Siebel and L. Zhou (2015). “Notch Receptor-Ligand Engagement Maintains Hematopoietic Stem Cell Quiescence and Niche Retention.” Stem Cells 33(7): 2280–2293.

Wang, W., G. Zimmerman, X. Huang, S. Yu, J. Myers, Y. Wang, S. Moreton, J. Nthale, A. Awadallah, R. Beck, W. Xin, D. Wald, A. Y. Huang and L. Zhou (2016). “Aberrant Notch Signaling in the Bone Marrow Microenvironment of Acute Lymphoid Leukemia Suppresses Osteoblast-Mediated Support of Hematopoietic Niche Function.” Cancer Res 76(6): 1641–1652.

Warr, M. R., E. M. Pietras and E. Passegué (2011). “Mechanisms controlling hematopoietic stem cell functions during normal hematopoiesis and hematological malignancies.” Wiley Interdiscip Rev Syst Biol Med 3(6): 681–701.

Wei, K., I. Korsunsky, J. L. Marshall, A. Gao, G. F. M. Watts, T. Major, A. P. Croft, J. Watts, P. E. Blazar, J. K. Lange, T. S. Thornhill, A. Filer, K. Raza, L. T. Donlin, C. W. Siebel, C. D. Buckley, S. Raychaudhuri and M. B. Brenner (2020). “Notch signalling drives synovial fibroblast identity and arthritis pathology.” Nature 582(7811): 259–264.

Wu, T., E. Hu, S. Xu, M. Chen, P. Guo, Z. Dai, T. Feng, L. Zhou, W. Tang, L. Zhan, X. Fu, S. Liu, X. Bo and G. Yu (2021). “clusterProfiler 4.0: A universal enrichment tool for interpreting omics data.” The Innovation 2(3): 100141.

Xiang, H., Y. Pan, M. A. Sze, M. Wlodarska, L. Li, K. A. van de Mark, H. Qamar, C. J. Moure, D. E. Linn, J. Hai, Y. Huo, J. Clarke, T. G. Tan, S. Ho, K. W. Teng, M. N. Ramli, M. Nebozhyn, C. Zhang, J. Barlow, C. E. Gustafson, S. Gornisiewicz, T. P. Albertson, S. L. Korle, R. Bueno, L. Y. Moy, E. H. Vollmann, D. Y. Chiang, P. E. Brandish and A. Loboda (2024). “Single-Cell Analysis Identifies NOTCH3-Mediated Interactions between Stromal Cells That Promote Microenvironment Remodeling and Invasion in Lung Adenocarcinoma.” Cancer Res 84(9): 1410–1425.

Yamamoto, W., E. Ogusa, K. Matsumoto, A. Maruta, Y. Ishigatsubo and H. Kanamori (2014). “Lymphocyte recovery on day 100 after allogeneic hematopoietic stem cell transplant predicts non-relapse mortality in patients with acute leukemia or myelodysplastic syndrome.” Leuk Lymphoma 55(5): 1113–1118.

Yang, S., S. E. Corbett, Y. Koga, Z. Wang, W. E. Johnson, M. Yajima and J. D. Campbell (2020). “Decontamination of ambient RNA in single-cell RNA-seq with DecontX.” Genome Biology 21(1): 57.

Yao, D., Y. Huang, X. Huang, W. Wang, Q. Yan, L. Wei, W. Xin, S. Gerson, P. Stanley, J. B. Lowe and L. Zhou (2011). “Protein O-fucosyltransferase 1 (Pofut1) regulates lymphoid and myeloid homeostasis through modulation of Notch receptor ligand interactions.” Blood 117(21): 5652–5662.

Yu, V. W., B. Saez, C. Cook, S. Lotinun, A. Pardo-Saganta, Y. H. Wang, S. Lymperi, F. Ferraro, M. H. Raaijmakers, J. Y. Wu, L. Zhou, J. Rajagopal, H. M. Kronenberg, R. Baron and D. T. Scadden (2015). “Specific bone cells produce DLL4 to generate thymus-seeding progenitors from bone marrow.” J Exp Med 212(5): 759–774.

Zhang, P., B. McGrath, S. Li, A. Frank, F. Zambito, J. Reinert, M. Gannon, K. Ma, K. McNaughton and D. R. Cavener (2002). “The PERK eukaryotic initiation factor 2 alpha kinase is required for the development of the skeletal system, postnatal growth, and the function and viability of the pancreas.” Mol Cell Biol 22(11): 3864–3874.

Zhong, L., L. Yao, R. J. Tower, Y. Wei, Z. Miao, J. Park, R. Shrestha, L. Wang, W. Yu, N. Holdreith, X. Huang, Y. Zhang, W. Tong, Y. Gong, J. Ahn, K. Susztak, N. Dyment, M. Li, F. Long, C. Chen, P. Seale and L. Qin (2020). “Single cell transcriptomics identifies a unique adipose lineage cell population that regulates bone marrow environment.” Elife 9.

