## Supplemental Figure for "Targeting Endothelial PERK Accelerates Lymphoid Regeneration by Enhancing DLL4-NOTCH3 Signaling at the Pre-B Niche"

**Supplementary figures**

**Supplemental Figure 1. Gating strategy for FACS analysis of bone marrow HSC and B progenitors, and impact of blocking IRE1α on HSC and B progenitors.**

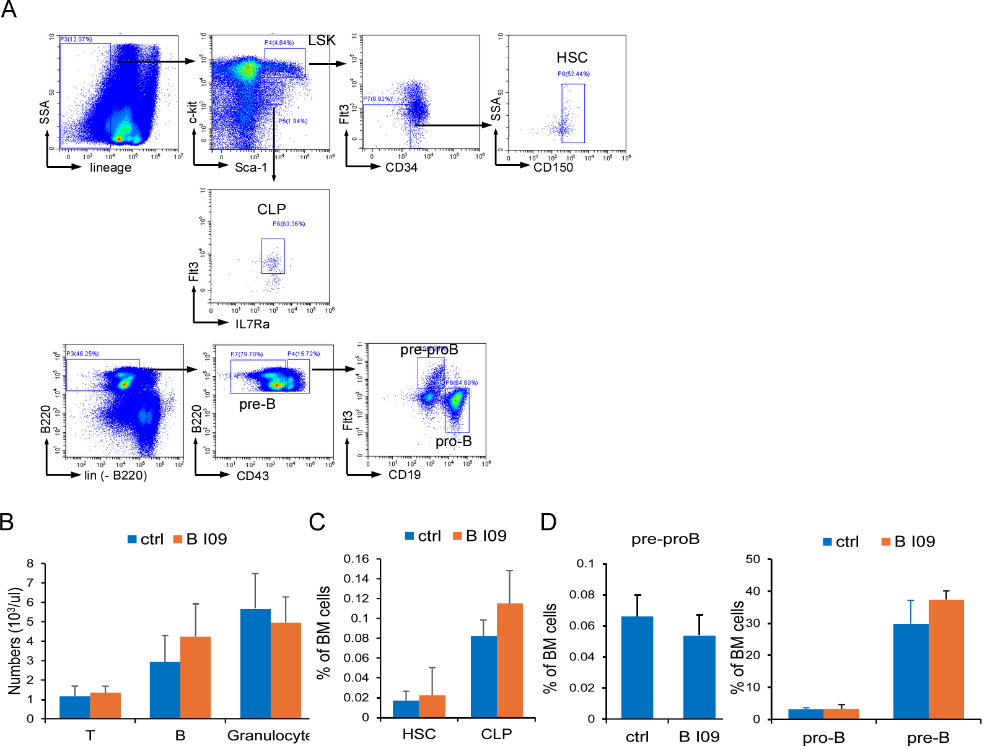

**(A)** Gating strategy for FACS analysis of BM HSC, CLP and B progenitors. For HSC and CLP, 7-AAD^-^Lin^-^ cell population was gated to define the expression of c-kit and Sca-1. LSKs (Lin^-^Sca-1^+^c-kit^+^) were then gated for HSC (LSK^+^CD34^-^Flt3^-^CD150^+^) analysis. Lin^-^Sca-1^low^c-kit^low^ population was gated for CLP (Lin^-^Sca-1^low^c-kit^low^Flt3^+^IL7R^+^) analysis. For B progenitor phenotyping, 7-AAD^-^Lin^-^ (CD11b, Gr1, CD4, CD8, Ter119, NK1.1) cell populations were gated to define the expression of B220 and CD43. Lin^-^B220^+^CD43^-^ were defined as pre-B while Lin^-^B220^+^CD43^+^ were gated for pre-proB and pro-B analysis (pre-proB: Lin^-^B220^+^CD43^+^Flt3^+^CD19^-^; pro-B: Lin^-^B220^+^CD43^+^Flt3^-^CD19^+^). **(B-D)** WT transplanted mice received IRE1α inhibitor BI09 (25 mg/kg) or control treatment for 5 days starting the day before irradiation. The treatment period was pre-determined to avoid toxicities associated with BI09 treatment longer than 5 days (data not shown). Peripheral blood B, T and granulocytes (B), BM HSC, CLP (C) and B progenitors (D) were determined on day 21. Results are presented as averages ± SD (n=7-9 in each group; data pooled from 3 experiments).

**Supplemental Figure 2. Blocking PERK boosted DLL4 expression and analysis of DKO and *Dll4*^iΔEC^ mice HSPCs.**

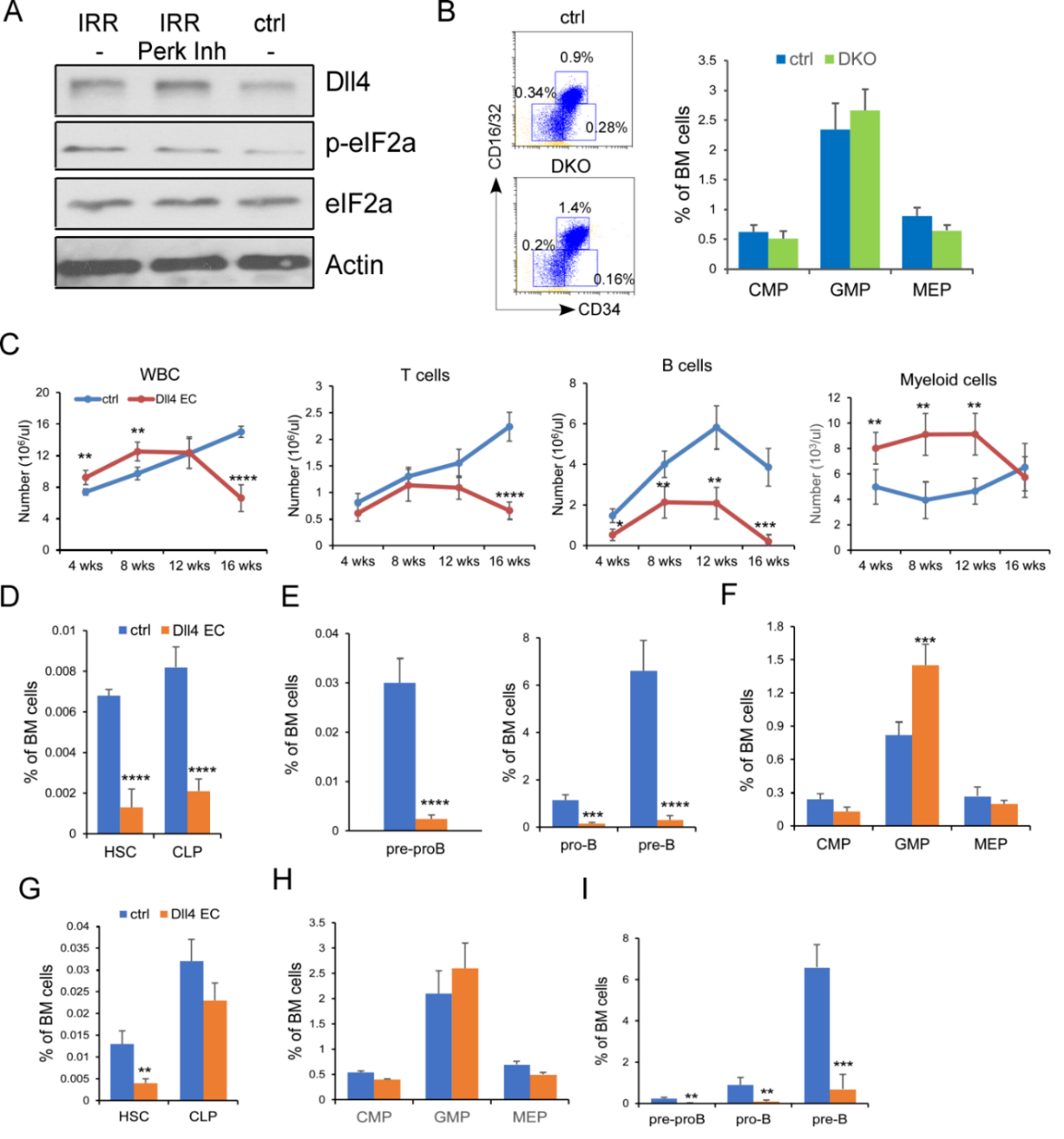

**(A)** Mouse BM endothelial cells (BMEC, Cellbiologics, C57-6221) were cultured in Complete Endothelial Cell Medium (Cellbiologics, M1168PF) at 37°C with 5% CO_2_. Cells undergoing irradiation received 850 cGy. For PERK inhibition, cells were treated with 1mM GSK2656157 for 24h and cells were harvested and subjected to Western blot. Shown was a representative blot of 3 similar experiments. **(B)** FACS was carried out on CytoFLEX. Representative FACS profile and frequencies of CMP/MEP/GMP in control and DKO mice at 1 month after transplantation. **(C)** Peripheral blood WBC, B cells, T cells, and granulocytes at 4, 8, 12, and 16 weeks after transplantation in WT and *Dll4*^iΔEC^ mice (n=7-8/group from 3 experiments). **(D-F)** Frequencies of HSC and CLP (D), B progenitors (E), and myeloid progenitors (F) were determined in control or *Dll4*^iΔEC^ recipient mice BM at 1 month after receiving WT donor BM cells in transplantation. Results were pooled from 3 experiments. **(G-I)** Frequencies of HSC and CLP (G), myeloid progenitors (CMP/MEP/GMP) (H), and B progenitors (I) at 4 months after transplantation in *Dll4*^iΔEC^ and control mice. Results in C-I are presented as averages ± SD. ***P*<0.01, *** *P*<0.001, **** *P*<0.0001

**Supplemental Figure 3. Cell clustering from scRNA-seq**.

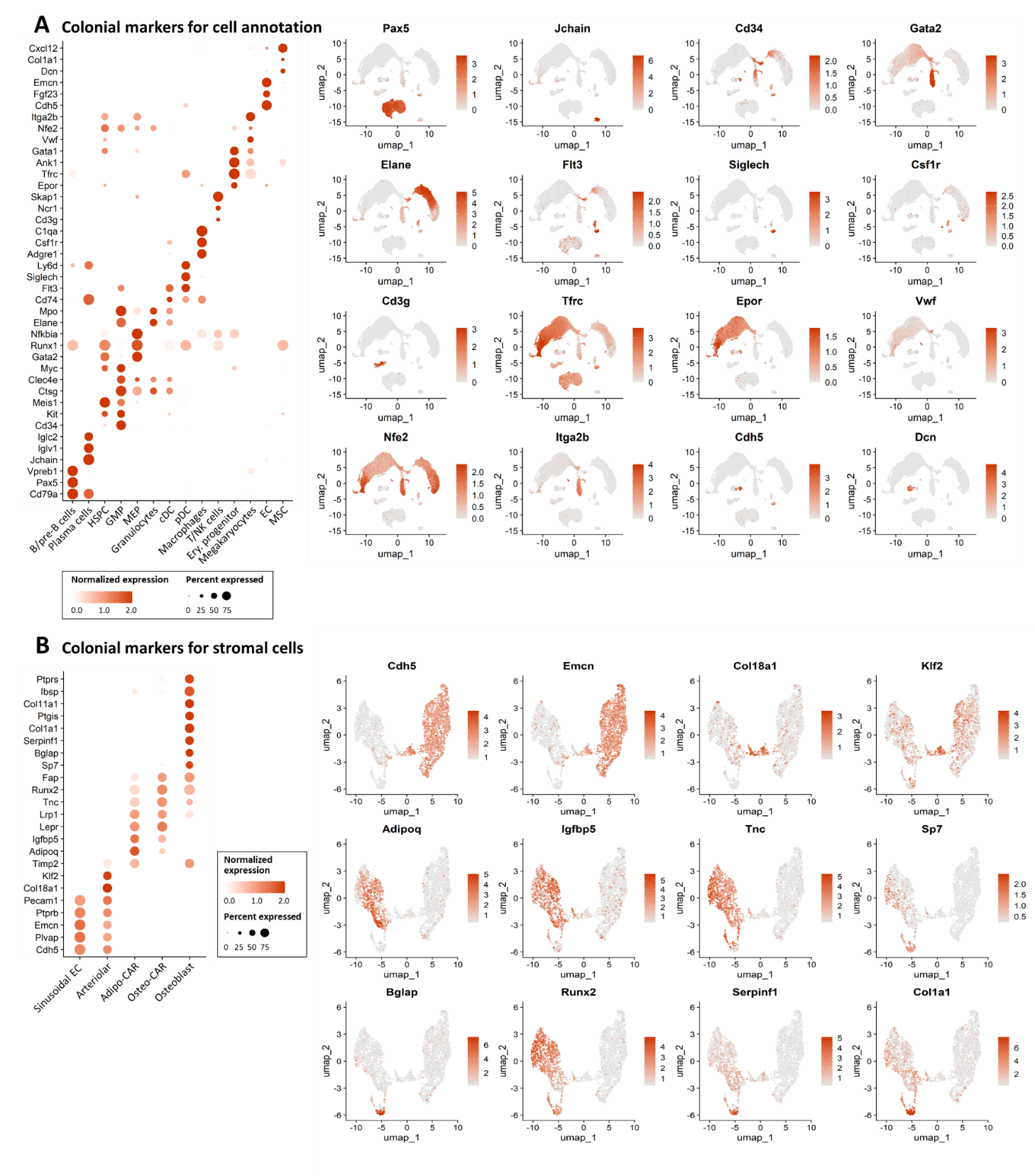

**(A-B)** Bubble plots and UMAP plots showing selection of cell type-specific markers across all major BM cell clusters (A) and stroma cell populations (B). The size of the dot indicates the fraction of cells expressing a particular marker, and the intensity of the color represents the level of mean expression.

**Supplemental Figure 4. Stem cell niche factors and interactions from scRNA-seq analysis.**

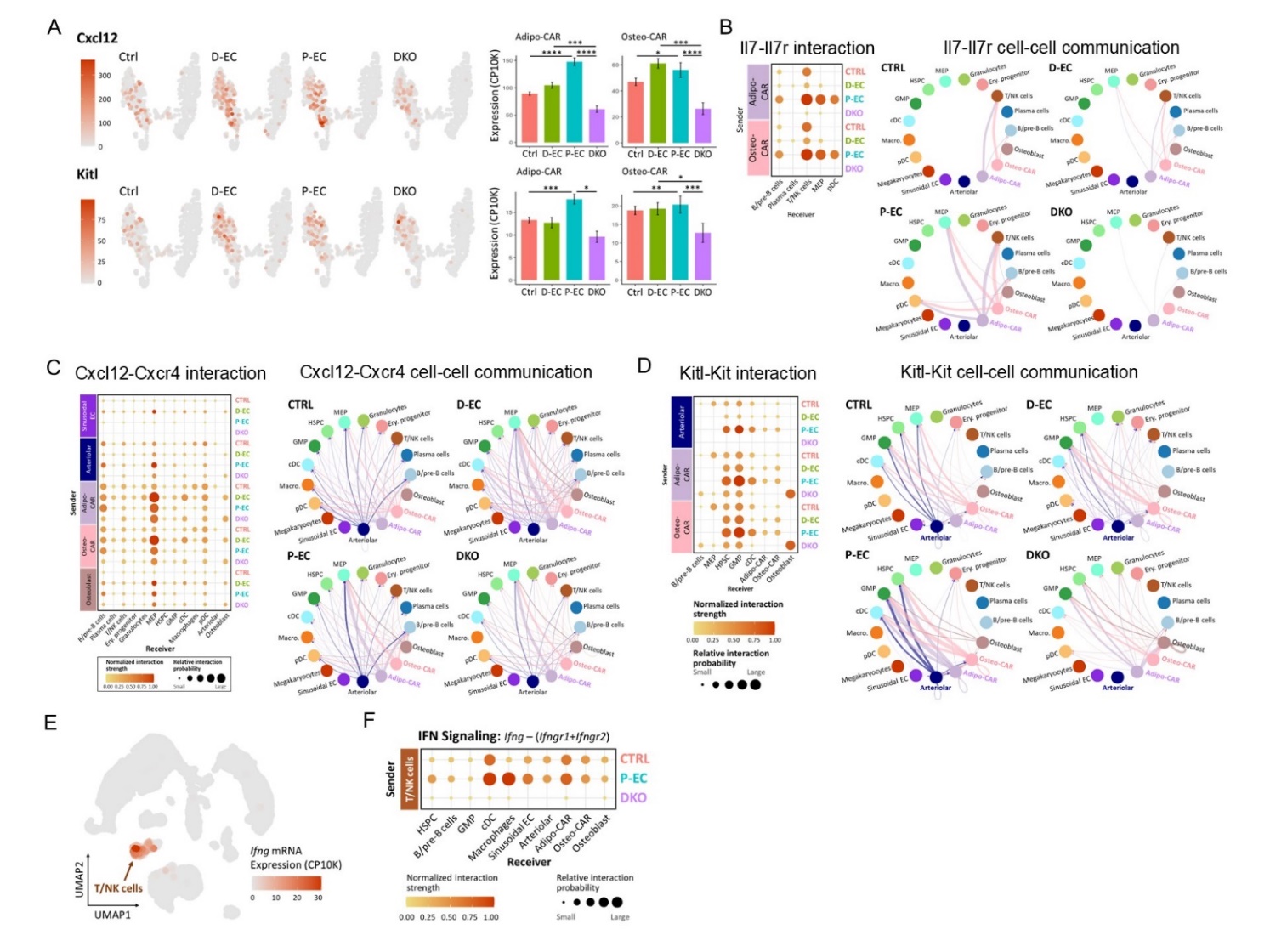

**(A)** UMAP and bar plots of *Cxcl12* and *Kitl* gene expression across all groups of mice. Statistical significances were first determined by Kruskal–Wallis test across four conditions (adjusted P<0.05), followed by post-hoc pairwise tests using MAST algorithm. **(B)** Bubble plots and circle plots of inferred Il7-Il7r signaling changes across conditions. Dot plot visualizes relative interaction strength of pathway signaling between adipo/osteo-CAR and their counterparts across different types of BM cells. **(C-D)** Bubble plot and circle plot of inferred Cxcl12-Cxcr4 (C) and SCF/KIT (D) signaling changes across conditions. **(E)** UMAP visualization of *Ifng* expression in T/NK cells. **(F)** Inferred IFN signaling (Ifng – Ifngr1+Ifngr2) changes across conditions. Dot plots visualize relative interaction strength of Ifng and receptor pairs between T/NK cells and their counterparts across treatment conditions.

**Supplemental Figure 5. Analysis of BM transplant and niche cytokine production in *Notch3^-/-^* mice.**
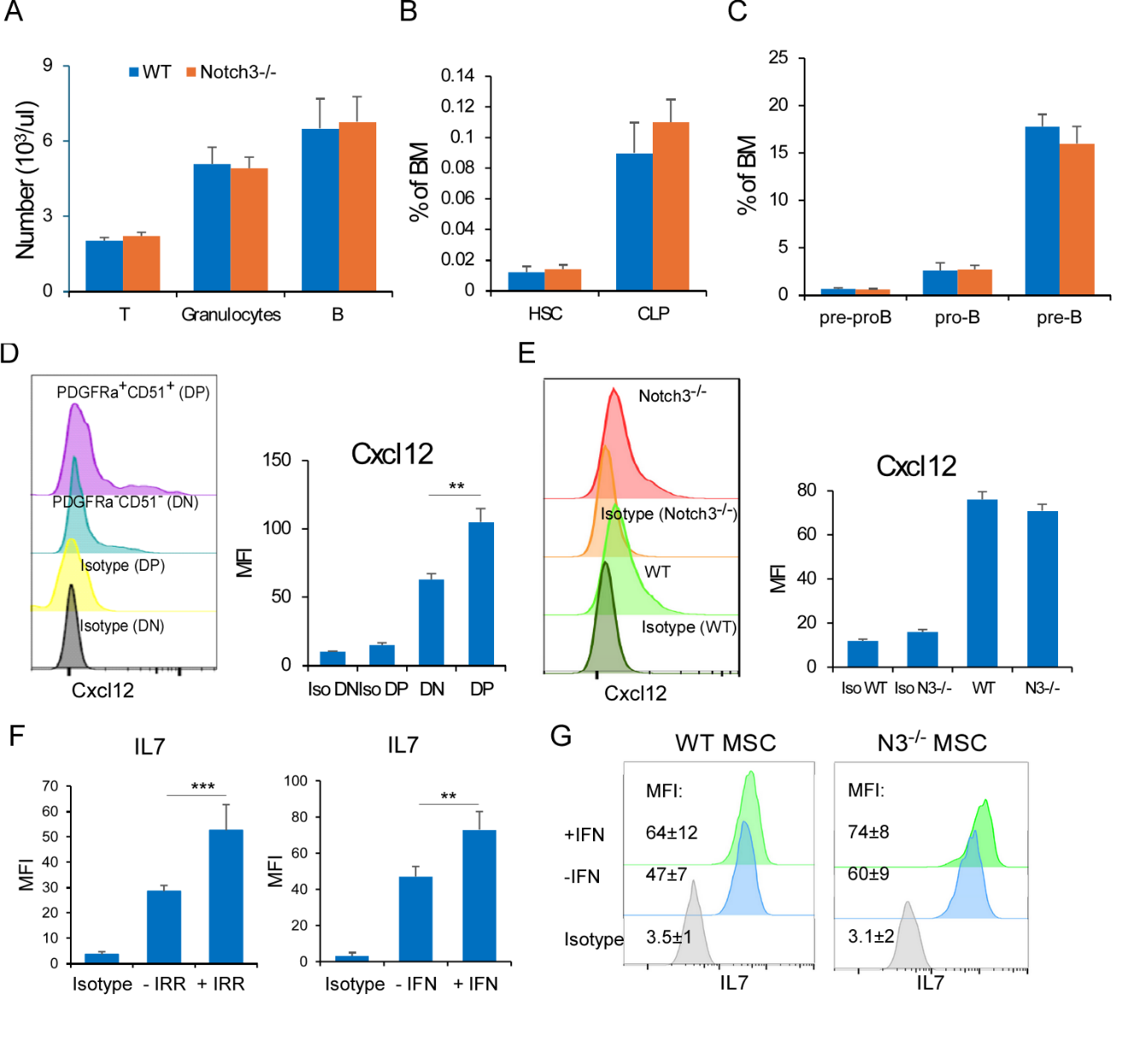

**(A)** BM transplantation was performed in WT mice (Ly5.1) receiving *Notch3^-/-^* or control WT donor cells (Ly5.2). Peripheral blood was analyzed for B cells, T cells, and granulocytes at 16 weeks after transplantation. **(B-C)** HSC, CLP and B progenitors were determined. Results from A-C were pooled from 2 experiments and presented as averages ± SD (n=6 / group). **(D)** Representative FACS profile and MFI of intracellular CXCL12 expression in mouse BM DP MSC (Lin^-^Ter119^-^CD31^-^PDGFRα^+^/CD51^+^) and DN MSC (Lin^-^Ter119^-^CD31^-^PDGFRα^-^/CD51^-^) (n=3/group). **(E)** Representative FACS profile and MFI of CXCL12 in WT and *Notch3^-/-^* BM DP MSC (n=3/group). **(F)** MFI of intracellular IL7 expression in BM DP MSCs of non-irradiated WT mice (-IRR) and irradiated WT mice (+IRR) on day 8 after receiving 550 cGy irradiation (left panel). In vitro cultured MSCs were briefly expanded, stimulated with IFNβ (1500 IU/ml) for 4 hrs. MFI of intracellular IL7 expression in DP MSCs was determined from 3 similar experiments (n=5/condition) (right panel). **(G)** FACS profile and MFI of IL7 expression in briefly expanded WT and *Notch3^-/-^* MSCs before and after receiving IFNβ (1500 IU/ml) for 4 hrs. Shown is one representative profile from 3 similar experiments.

**Supplemental Figure 6. Analysis of hematopoietic and vascular regeneration in *Jag1*^iΔEC^ mice.**

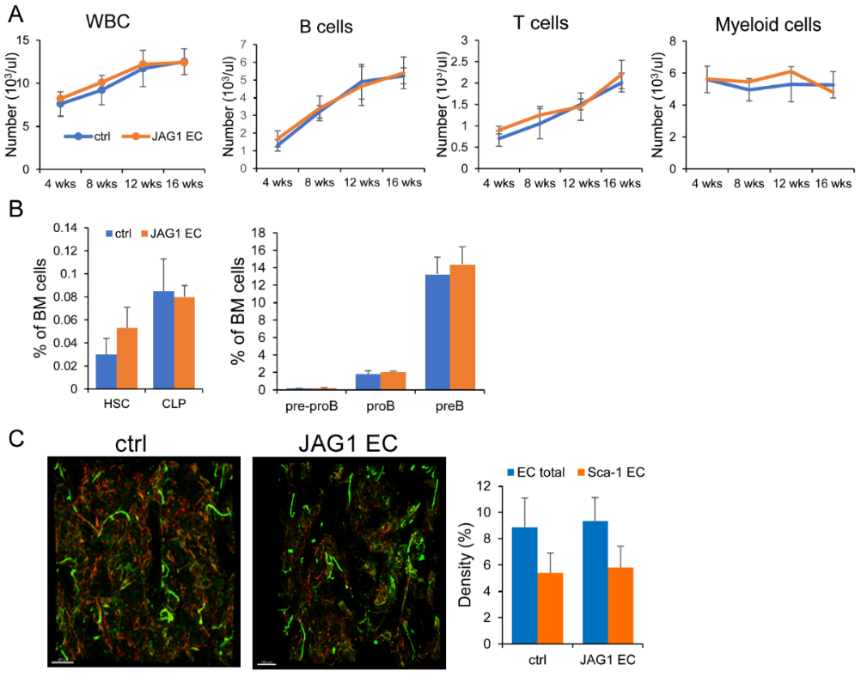

**(A)** BM transplantation was performed in *Jag1*^iΔEC^ and control mice as described for *Dll4*^iΔEC^ mice. Peripheral blood was analyzed for WBC counts, B cells, T cells, and granulocytes at 4, 8, 12, and 16 weeks after transplantation (n=6-8/group). **(B)** HSC, CLP and B progenitors were determined. Results from A-B were pooled from 2 experiments and presented as averages ± SD (n=6-8/group). **(C)** Whole-mount immunostaining of BM vascular bed with Alexa Fluor 647 anti-CD31/anti-CD144 and FITC-anti-Sca-1. Representative images were generated in Imaris based on the corresponding fluorescent signals (left). The percentage of total CD31^+^/CD144^+^ volume (EC total) and CD31^+^Sca-1^+^ arteriole volume (Sca-1 EC) were determined (right). Results are presented as averages ± SD (n=5 in each group).
